# Wistar rats and C57BL/6J mice differ in their motivation to seek social interaction versus food in the Social versus Food Preference Test

**DOI:** 10.1101/2020.06.04.134437

**Authors:** C.J. Reppucci, L.A. Brown, A.Q. Chambers, A.H. Veenema

## Abstract

Here we characterized the Social versus Food Preference Test, a behavioral paradigm designed to investigate the competition between the choice to seek social interaction versus the choice to seek food. We assessed how this competition was modulated by internal cues (social isolation, food deprivation), external cues (time-of-testing, stimulus salience), sex (males, females), age (adolescents, adults), and rodent model (Wistar rats, C57BL/6J mice). We found that changes in stimulus preference in response to the internal and external cue manipulations were similar across cohorts. Specifically, social over food preference scores were reduced by food deprivation and social familiarly in Wistar rats and C57BL/6J mice of both sexes. Interestingly, the degree of food deprivation-induced changes in stimulus investigation patterns were greater in adolescents compared to adults in Wistar rats and C57BL/6J mice. Strikingly, baseline stimulus preference and investigation times varied greatly between rodent models: across manipulations, Wistar rats were generally more social-preferring and C57BL/6J mice were generally more food-preferring. Adolescent Wistar rats spent more time investigating the social and food stimuli than adult Wistar rats, while adolescent and adult C57BL/6J mice investigated the stimuli a similar amount. Neither social isolation nor time-of-testing altered behavior in the Social versus Food Preference Test. Together, our results indicate that the Social versus Food Preference Test is a flexible behavioral paradigm suitable for future interrogations of the peripheral and central systems that can coordinate the expression of stimulus preference related to multiple motivated behaviors.

**HIGHLIGHTS:**

- Rats prefer social over food when sated, and this is attenuated by food deprivation.
- Mice have no preference when sated, and prefer food over social when food-deprived.
- Rats prefer a familiar social stimulus or a novel social stimulus over food.
- Mice prefer food over a familiar social stimulus.
- Adolescent rats investigate social and food stimuli longer than adult rats.

**GRAPHICAL ABSTRACT:** 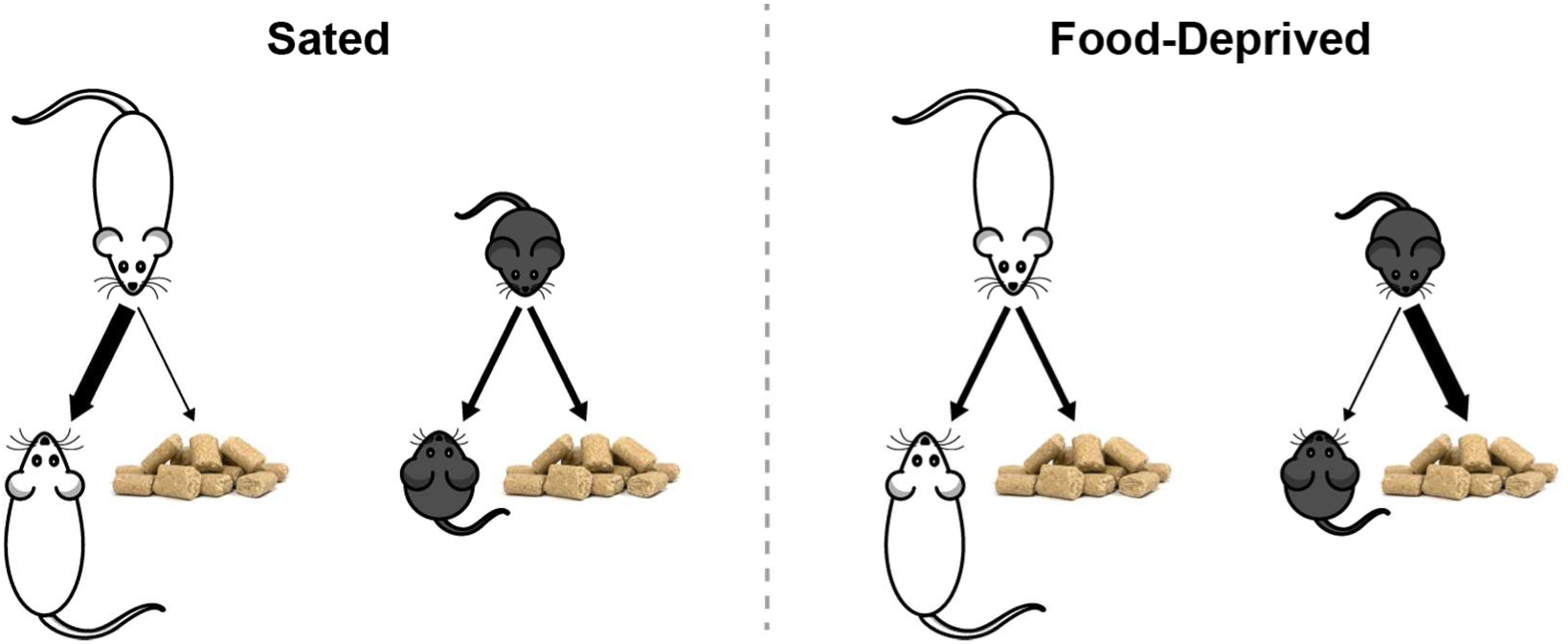

## 1 INTRODUCTION

Behavior is influenced by a combination of internal and external cues, with the expression of the appropriate behavior dependent on an individual’s current motivational state and the presence of stimuli in their surrounding environment [1, 2]. Thus far, most laboratory studies have focused on uncovering the peripheral and central systems that regulate the expression of a single behavior or the expression of a suite of behaviors associated with a single motivational state. In daily life, however, an individual can be experiencing, at the same moment, multiple motivational states with multiple choices of how to act. Yet, the direct assessment of the roles of peripheral and central systems in coordinating motivated behavioral choice is largely understudied [3-6]. This may be due to a lack of behavioral tests that are suitable for such investigations. Here, we characterized a recently developed behavioral paradigm [3], hereafter called the Social versus Food Preference Test, to test the competition between the choice to seek social interaction versus the choice to seek food.

To determine how internal cues modulate the competition between the choice to seek social interaction versus the choice to seek food, we altered the motivational states of subjects by exposing them to acute social isolation and/or acute food deprivation prior to exposure to the Social versus Food Preference Test. To examine whether the effects of these manipulations were similar between the sexes, stable across the lifespan, and comparable between commonly used laboratory rodent models, experiments were conducted with adolescent and adult Wistar rats and C57BL/6J mice of both sexes (Experiments 1 and 3). For all subjects, we predicted that social isolation biases preference towards the social stimulus, and that food deprivation biases preference towards the food stimulus.

To determine how external cues modulate the competition between the choice to seek social interaction versus the choice to seek food, we tested whether the time-of-testing influences food deprivation-induced changes in social versus food preference in adolescent male and female Wistar rats (Experiment 2), and whether the saliency of the social stimulus alters social versus food preference in adolescent Wistar rats and C57BL/6J mice of both sexes (Experiment 4). We predicted that the time-of-testing alters locomotor activity [7] but not social versus food preference, and that stimulus preference is more biased toward the social stimulus when the social stimulus is novel compared to when the social stimulus is familiar [8-10].

Lastly, to assess general sociability in C57BL/6J mice [9], we tested the preference of adolescent and adult mice of both sexes to investigate a social stimulus versus an empty corral (Experiment 5). In sum, the series of experiments presented in this paper aimed to characterize a flexible behavioral paradigm suitable for future interrogations of the peripheral and central systems that coordinate the choice to seek social interaction versus the choice to seek food.

## 2 MATERIALS AND METHODS

### 2.1 Animals

Male and female Wistar rats (Charles River Laboratories) were housed in single sex groups of two to four in standard rat cages (48 × 27 × 20 cm), and male and female C57BL/6J mice (Jackson Laboratories’ stock 000664 or Charles River Laboratories) were housed in single sex groups of two to four in standard mouse cages (29 × 19 × 13 cm). Rats and mice were housed in separate colony rooms within the vivarium, and all animals were maintained under standard laboratory conditions (12 hr light/dark cycle; water *ad libitum*; food *ad libitum* except as described in **Section 2.3**). All housing and testing was in accordance with the National Institute of Health *Guidelines for Care and Use of Laboratory Animals* and the Michigan State University Institutional Animal Care and Use Committee. Animals were acclimated to the colony rooms for at least 72 hrs prior to the start of daily handling or any experimental procedures. All animals were gonadally intact, but estrous cycle was not monitored.

### 2.2 Social versus Food Preference Test

The Social versus Food Preference Test was used to assess the preference of rats and mice to investigate a social stimulus (age-, sex-, and species-matched conspecific) versus a food stimulus (standard laboratory chow; Teklad Irradiated 22/5 Rodent Diet, 8940). This test was based on our previously developed social novelty preference test in rats [8], and a social interaction assay used to determine the effects of hunger signals on social interest in mice [3]. Social versus food preference was tested using a 3-chambered apparatus, where the social stimulus and the food stimulus were located on opposite ends. Two sizes of this apparatus were custom-constructed (**Fig 1**), one for rats (Scientific Instrumental and Machining Services, Boston College) and one for mice (Physics and Astronomy Machine Shop, Michigan State University), and each testing apparatus was located in its respective colony room within the vivarium. The exterior of the apparatus was composed of Plexiglas (rats) or PVC (mice), and each chamber (rats: 40 cm × 40 cm × 27 cm; mice: 30 cm × 30 cm × 20 cm) was separated by a translucent Plexiglas (rats) or acrylic (mice) partition with an opening (rats: 10 cm × 10.2 cm; mice: 5 cm × 5 cm) to allow passage between chambers. Stimuli were placed in corrals, which allowed for olfactory, visual, and auditory contact, but restricted tactile contact by the experimental subject (food pellets were moved away from accessible edges to prevent consumption). For rats, rectangular corrals (18 cm × W 10 cm D × 21 H cm) were composed of a solid translucent Plexiglas top/bottom/back and translucent Plexiglas bars (0.6 cm diameter, spaced 1.75 cm apart) on the other three sides. For mice, cylindrical corrals (8.5 cm ID, 10.5 cm OD × 17cm H) were composed of solid translucent Plexiglas top/bottom connected by translucent Plexiglas bars (0.6 cm diameter, spaced 1.5 cm apart). The apparatus was cleaned with 70% ethanol and corrals were cleaned with dilute cleaning solution at the start and end of each day, as well as between subjects.

**Fig 1.**
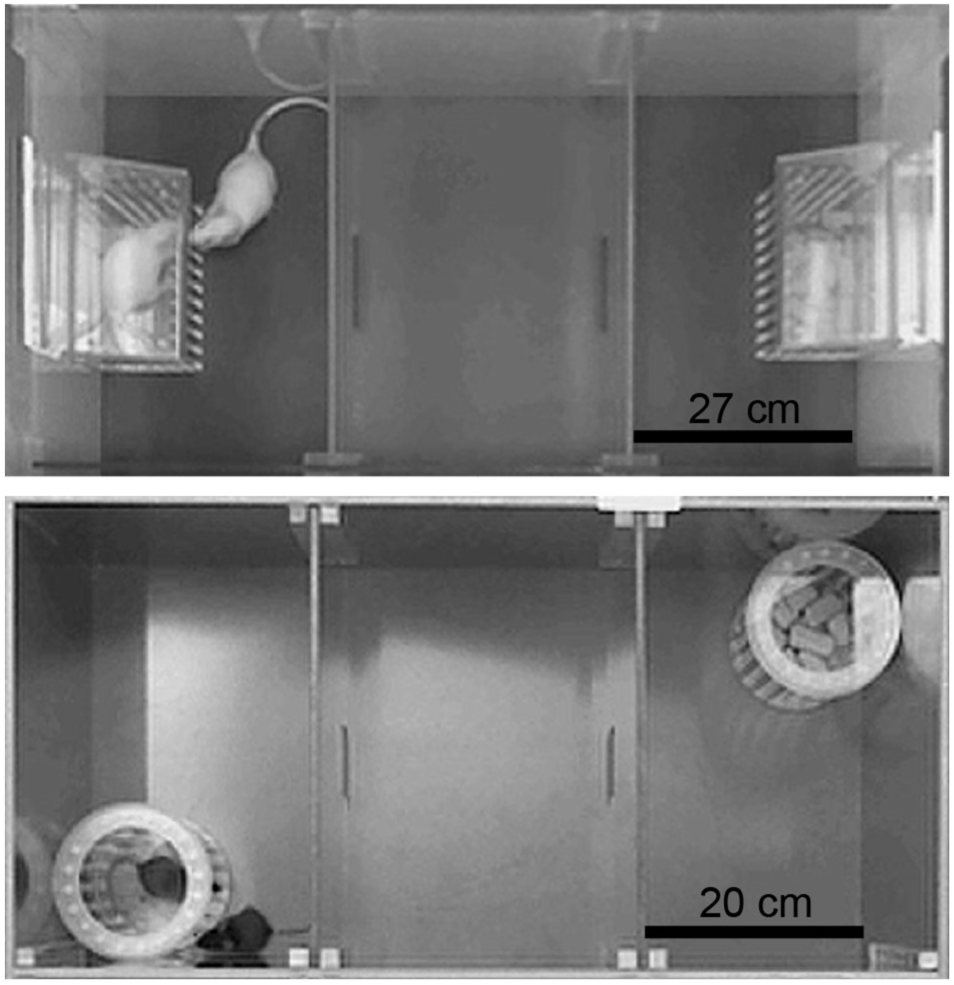
Social versus Food Preference Test. Rats (top) and mice (bottom) were placed into the center of a 3-chambered apparatus and then allowed to freely investigate a social stimulus and a food stimulus, which were placed in corrals located on opposite ends, for a period of 10 min.

All subjects were habituated to the testing procedures the day prior to their first test. During habituation, experimental animals were placed into the center chamber and allowed to freely explore the apparatus and investigate empty corrals for 10 min before being returned to their homecage. In separate trials, stimulus animals were habituated to confinement within a corral for 10 min before being returned to their homecage. There were no significant chamber preferences during habituation in any experiment (data not shown), and data from habituation sessions were not used in any other statistical analyses.

During testing, the experimental subject was placed into the center chamber and allowed to freely explore the apparatus and investigate the corralled stimuli for 10 min. To reduce the number of animals used, when the social stimulus was novel it was used twice per day (exposed to two different experimental subjects in non-successive tests); food pellets were replaced between subjects. The location of the social and food stimuli (i.e., left chamber or right chamber) was pseudorandom and counterbalanced between subjects each day and within subjects across test days (when applicable). A webcam (Logitech HD Pro C910) was attached to the ceiling and connected to a PC computer in an adjoining room to record the sessions. Experimenters, who were unaware of sex and testing conditions, scored recorded videos using Solomon Coder (https://solomon.andraspeter.com/) to quantify the amount of time the experimental subjects spent investigating each of the two stimuli, and these values were then summed to calculate total stimulus investigation time. Investigation was defined as when the subject was actively engaged with the corral (e.g., sticking nose between bars, pawing, sniffing) with its attention directed towards the stimulus inside of the corral as indicated by head position/gaze orientation. To obtain a measure of stimulus preference, the percent of time the subject investigated the social stimulus [(time spent investigating social stimulus/total stimulus investigation time)*100] was calculated (“social over food preference score”). Values > 50% indicate that subjects spent more time investigating the social stimulus, and values < 50% indicate that subjects spent more time investigating the food stimulus. AnyMaze (Stoelting) was used to quantify measures of locomotor activity (i.e., middle chamber entries, distance traveled). In instances where AnyMaze failed to track the experimental subject, experimenters manually scored videos for middle chamber entries and these subjects were removed from the distance traveled analyses (Experiment 1a: n = 5 females; Experiment 1b: n = 1 male, n = 4 females, Experiment 2: n = 2 females, Experiment 3a: n = 2 males, n = 2 females, Experiment 3b: n = 1 female, Experiment 4b: n = 1 female).

### 2.3 Experimental Procedures

#### 2.3.1 Experiment 1: The effects of social isolation and food deprivation on social versus food preference in rats

To establish the Social versus Food Preference Test and characterize how manipulating the internal motivational drives for social interaction-seeking and food-seeking behavior affects behavior in this test, we tested how subjects’ preference to investigate a social stimulus (novel age- and sex-matched conspecific) versus a food stimulus (standard laboratory chow) was modulated by social isolation and hunger. In Experiment 1a the subjects were adolescent (39-46 day old) Wistar rats (8 males/6 females; 1 male was subsequently removed from all analyses due to escaping the testing apparatus), and in Experiment 1b the subjects were a separate cohort of adult (13-14 week old) Wistar rats (8 males/8 females). In both experiments, subjects were first habituated to the testing apparatus as described above, and then tested in the Social versus food Preference Test at zeitgeber time (ZT) 12 on four occasions each 48 hrs apart using a within-subjects 2 × 2 counterbalanced design (pair-housed/socially isolated x sated/food-deprived). The length of social isolation and/or food deprivation was 24 hrs, and experimental subjects were exposed to a different unfamiliar social stimulus during each test. Subjects’ body weights during this 24 hr period were monitored as a proxy physiological measure of hunger. Subjects were returned to their homecage immediately following the Social versus Food Preference Test, and *ad lib* chow consumption during the subsequent 30 min was monitored to behaviorally assess hunger. As such, when subjects were tested under pair-housed conditions consumption was measured as the amount eaten by the pair, and when subjects were tested under isolated conditions consumption was the amount eaten by each individual. Percent change in body weight [((end weight-start weight)/start weight)*100] and consumption as a percent of body weight [(total grams consumed/total grams body weight)*100] were calculated to normalize the data across experiments and account for baseline sex differences in body weight which could also influence consumption potentially due to differences in gastric capacity [11] or other factors [12]. Lastly, food was removed from the stimulus animals’ homecages 2 hrs prior to the start of the testing each day to reduce the amount of food-related sensory cues present on the social stimuli [as described in: 3]. At the end of each test session, animals were returned to their original pair-housing conditions with *ad lib* food access.

#### 2.3.2 Experiment 2: Does the time-of-testing modulate food deprivation-driven changes in social versus food preference in adolescent rats?

To determine whether the stimulus preference and investigation patterns observed in Experiment 1a could be explained by the time-of-testing, we tested how subjects’ preferences to investigate the social stimulus (novel age- and sex-matched conspecific) versus the food stimulus (standard laboratory chow) were modulated by food deprivation and the time-of testing. Specifically, we compared behavior during the start of the dark phase (ZT12) to behavior during the middle of the light phase (ZT7). Adolescent (39-46 day old) Wistar rats (6 males/8 females) were first habituated to the testing apparatus and then tested on four occasions each 48 hrs apart using a within-subjects 2 × 2 counterbalanced design (sated/food-deprived x dark phase/light phase). All other experimental procedures were as described in Experiment 1a, with the exception that experimental animals were single-housed for the entire duration of this experiment (i.e., 8 days of social isolation in Experiment 2 versus acute 24 hr social isolation in Experiment 1a), and thus all consumption measures represented consumption by individual subjects.

#### 2.3.3 Experiment 3: The effects of social isolation and food deprivation on social versus food preference in mice

To determine whether the stimulus preference and investigation patterns observed in Experiment 1 are conserved across commonly used laboratory rodent models, we repeated this experiment in mice. In Experiment 3a the subjects were adolescent (37-44 day old) C57BL/6J mice (8 males/8 females; 1 female was subsequently removed from all analyses for spending, on average, 3 min of each test behind a corral and out-of-view of the experimenter), and in Experiment 3b the subjects were a separate cohort of adult (13-14 week old) C57BL/6J mice (8 males/8 females; 1 female was subsequently removed from all analyses for failure to explore all three chambers during habituation or test one). The following changes were implemented: adolescent subjects were tested 2 days younger than in Experiment 1a to account for the mildly faster development of mice compared to rats [13], testing started at ZT7 (based on outcomes of Experiment 2), the length of social isolation/food deprivation was reduced to 18 hrs following pilot testing to identify conditions that ensured mice would not lose more than 15% of their body weight, and post-test food consumption was conducted individually in clean cages to allow for within-subjects comparison of consumption by subjects tested under pair-housed and isolated conditions (after the consumption test, mice were re-housed in their homecages).

#### 2.3.4 Experiment 4: Modulation of social versus food preference by social salience in sated adolescent rats and mice

To determine whether stimulus preference and investigation patterns is modulated by the salience of external stimuli, we tested subjects’ preferences to investigate a novel (age-, sex-, and species-matched conspecific) or familiar (cagemate) social stimulus versus the food stimulus (standard laboratory chow) in subjects that were maintained under pair-housing and *ad lib* feeding conditions. Subjects were first habituated to the testing apparatus and then using a within-subjects counterbalanced design each subject was tested on two occasions 48 hrs apart, with testing starting at ZT7. In Experiment 4a, the subjects were adolescent (39-42 day old) Wistar rats (9 males/8 females), and in Experiment 4b the subjects were adolescent (37-40 day old) C57BL/6J mice (8 males/8 females). Cagemates were not used as novel social stimuli for other experimental subjects. Because subjects were tested under sated conditions and cagemates sometimes served as stimulus animals, food was not removed from the homecages of stimulus animals prior to testing.

#### 2.3.5 Experiment 5: Do sated mice prefer to investigate a novel social stimulus over an empty corral?

To assess general sociability in C57BL/6J mice, we tested subjects’ preferences to investigate a social stimulus (novel age-, sex-, and species-matched conspecific) versus an empty corral. Subject mice were maintained under pair-housing and *ad lib* feeding conditions, habituated to the testing apparatus, and tested the following day starting at ZT7. In Experiment 5a the subjects were adolescent (44 day old) C57BL/6J mice (4 male/4 female; formerly stimulus animals in Experiment 3a), and in Experiment 5b the subjects were a separate cohort of adult (15 week old) C57BL/6J mice (4 male/4 female; formerly stimulus animals in Experiment 3b). As in Experiments 1-3, food was removed from the homecages of the stimulus animals 2 hrs prior to the start of testing.

### 2.4 Statistical analysis

Mixed-model [sex (male, female; between-subjects factor) x hunger condition (food-deprived, sated; within-subjects factor) x housing condition (socially isolated, pair-housed; within-subjects factor)] ANOVAs were used to assess the effects of sex, food deprivation, and social isolation on stimulus preference, stimulus investigation, locomotor activity, and body weight measures in Experiments 1a, 1b, 3a, and 3b, as well as consumption measures in Experiments 3a and 3b. Since consumption was assessed under pair-housed conditions in Experiments 1a and 1b (see **Section 2.3.1**), separate mixed-model [sex (male, female; between-subjects factor) x hunger condition (food-deprived, sated; within-subjects factor)] ANOVAs were used to evaluate consumption under social isolation and pair-housing. Mixed-model [sex (male, females; between-subjects factor) x hunger condition (food-deprived, sated; within-subjects factor x time-of-testing (dark phase, light phase; within-subjects factor)] ANOVAs were used to assess the effects of sex, food deprivation, and time-of-testing for all measures in Experiment 2. Mixed-model [sex (male, females; between-subjects factor) x social saliency (novel, cagemate; within-subjects factor)] ANOVAs were used to assess the effects of sex and social salience for all measures in Experiments 4a and 4b. Mixed-model ANOVAs [sex (male, female; between-subjects factor) x stimulus (social corral, empty corral; within-subjects factor)] investigated sex differences in corral investigation times in Experiments 5a and 5b. Independent samples t-Tests were used to assess sex differences in stimulus preference and locomotor activity in Experiments 5a and 5b. When significant interactions were found in the ANOVAs, Bonferroni *post hoc* pairwise comparisons were conducted to clarify the effects. For all experiments, one-sample t-Tests with a reference value of 50% were used to evaluate stimulus preference.

Normality was assessed by Shapiro-Wilk testing of the standardized residuals, sphericity by Mauchly’s Test, and equality of variances by Levene’s Test. All data were analyzed using IBM SPSS Statistics 24-26, and statistical significance was set at *p* < 0.05. Cohen’s d (*d*) effect sizes were manually computed for all t-Tests, and partial eta squared (η^2^) effect sizes were computed in SPSS for all ANOVAs. Graphs were produced in Microsoft Excel using custom templates and those from [14], and then edited in Adobe Illustrator CC.

## RESULTS

### 3.1 Experiment 1a: Social versus food preference was altered by food deprivation, but not social isolation in adolescent rats

Adolescent rats had significantly lower social over food preference scores when tested under food-deprived compared to sated conditions; neither housing condition nor sex altered stimulus preference (**Table 1, Fig 2A, B**). Adolescent rats had a strong social preference under sated conditions (pair-housed: *t*_(12)_ = 20.0, *p* < 0.001, *d* = 5.55; socially isolated: *t*_(12)_ = 17.1, *p* < 0.001, *d* = 4.74), and no stimulus preference under food-deprived conditions (pair-housed: *t*_(12)_ = 1.40, *p* = 0.19, *d* = 0.39; socially isolated: *t*_(12)_ = 0.14, *p* = 0.89, *d* = 0.039; **Fig 2A, B**).

**Table 1.** ANOVA statistics and partial eta squared (η^2^) effect sizes for Experiment 1a: Social versus food preference was altered by food deprivation, but not social isolation in adolescent rats. Significant effects shown in **bold**, n.s. = none significant, n.a. = not applicable, BW = body weight.

|  | Sex | Hunger Condition | Housing Condition | Interactions |
| --- | --- | --- | --- | --- |
| Social over Food Preference [%] | $F_{(1,11)} = 1.95, p = 0.19, \eta^2 = 0.15$ | $F_{(1,11)} = \mathbf{58.5}, p < \mathbf{0.001}, \eta^2 = \mathbf{0.84}$ | $F_{(1,11)} = 1.36, p = 0.27, \eta^2 = 0.11$ | n.s. |
| Social Stimulus Investigation [sec] | $F_{(1,11)} = 0.84, p = 0.38, \eta^2 = 0.071$ | $F_{(1,11)} = \mathbf{9.83}, p = \mathbf{0.009}, \eta^2 = \mathbf{0.47}$ | $F_{(1,11)} = 0.00, p = 0.99, \eta^2 < 0.001$ | n.s. |
| Food Stimulus Investigation [sec] | $F_{(1,11)} = 2.77, p = 0.12, \eta^2 = 0.20$ | $F_{(1,11)} = \mathbf{88.6}, p < \mathbf{0.001}, \eta^2 = \mathbf{0.89}$ | $F_{(1,11)} = 3.05, p = 0.11, \eta^2 = 0.22$ | n.s. |
| Total Investigation [sec] | $F_{(1,11)} = 0.001, p = 0.98, \eta^2 < 0.001$ | $F_{(1,11)} = 1.18, p = 0.30, \eta^2 = 0.097$ | $F_{(1,11)} = 1.23, p = 0.29, \eta^2 = 0.10$ | n.s. |
| Middle Chamber Entries [#] | $F_{(1,11)} = 1.97, p = 0.19, \eta^2 = 0.15$ | $F_{(1,11)} = \mathbf{15.5}, p = \mathbf{0.002}, \eta^2 = \mathbf{0.59}$ | $F_{(1,11)} = 1.92, p = 0.19, \eta^2 = 0.15$ | n.s. |
| Body Weight [% change] | $F_{(1,11)} = 2.03, p = 0.18, \eta^2 = 0.16$ | $F_{(1,11)} = \mathbf{241}, p < \mathbf{0.001}, \eta^2 = \mathbf{0.96}$ | $F_{(1,11)} = \mathbf{11.7}, p = \mathbf{0.006}, \eta^2 = \mathbf{0.51}$ | n.s. |
| Consumption [% of BW] – Pair-Housed | $F_{(1,4)} = 0.53, p = 0.51, \eta^2 = 0.12$ | $F_{(1,4)} = \mathbf{111}, p < \mathbf{0.001}, \eta^2 = \mathbf{0.97}$ | n.a. | n.s. |
| Consumption [% of BW] – Socially Isolated | $F_{(1,11)} = 1.94, p = 0.19, \eta^2 = 0.15$ | $F_{(1,11)} = \mathbf{31.3}, p < \mathbf{0.001}, \eta^2 = \mathbf{0.74}$ | n.a. | n.s. |

**Fig 2.**
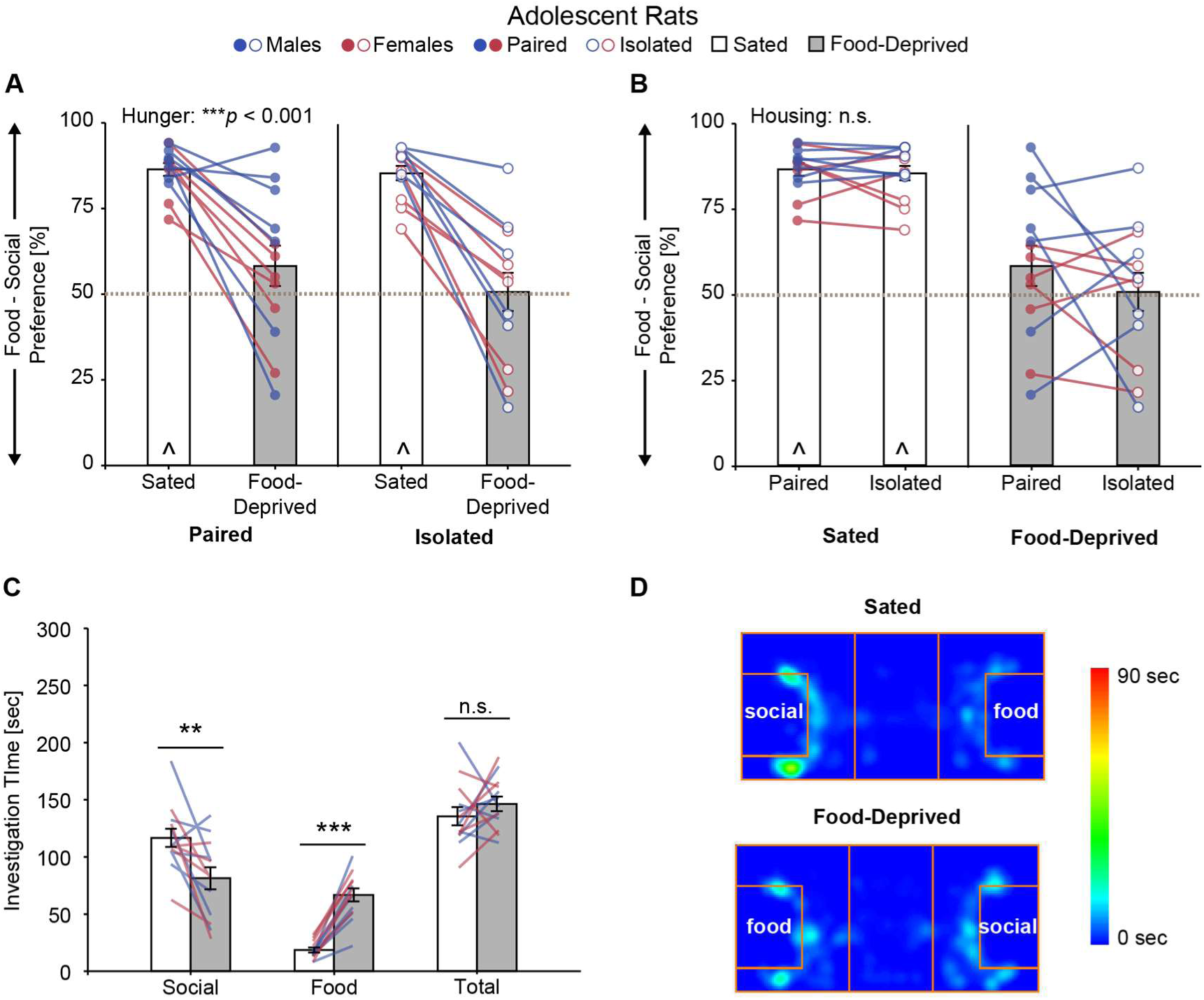
Experiment 1a. Food deprivation abolished preference for the social stimulus in adolescent rats (**A**), while social isolation had no effect on stimulus preference (**B**). Food deprivation decreased investigation of the social stimulus, and robustly increased investigation of the food stimulus (**C**). Representative heat maps of activity from one subject (**D**). Bar graphs (mean ± SEM) collapsed across sex for **A** and **B** (same data replotted to illustrate main effects), and across sex and housing condition for **C**; individual data collapsed across housing condition in **C**; * *p* < 0.05, ** *p* < 0.01, *** *p* < 0.001 mixed-model ANOVA; ^ p < 0.05, one-sample t-Test from 50% (gray dashed line); n.s. = not significant; n = 7 males, n = 6 females.

The observed changes in stimulus preference were due to decreased time spent investigating the social stimulus and increased time spent investigating the food stimulus when adolescent rats were food-deprived compared to when they were sated (**Table 1, Fig 2C**). Despite food deprivation-induced changes in the time spent investigating each stimulus, there was no net change in total (social + food) investigation time as a result of food deprivation (**Table 1, Fig 2C**). Stimulus investigation times were unaffected by housing condition or sex (**Table 1**).

Locomotor activity, as measured by middle chamber entries, was decreased under food-deprived compared to sated conditions and unaffected by housing condition or sex in adolescent rats (**Tables 1, 2**). Distance traveled was not assessed due to technical difficulties tracking the female cohort (see **Section 2.2**).

**Table 2.**
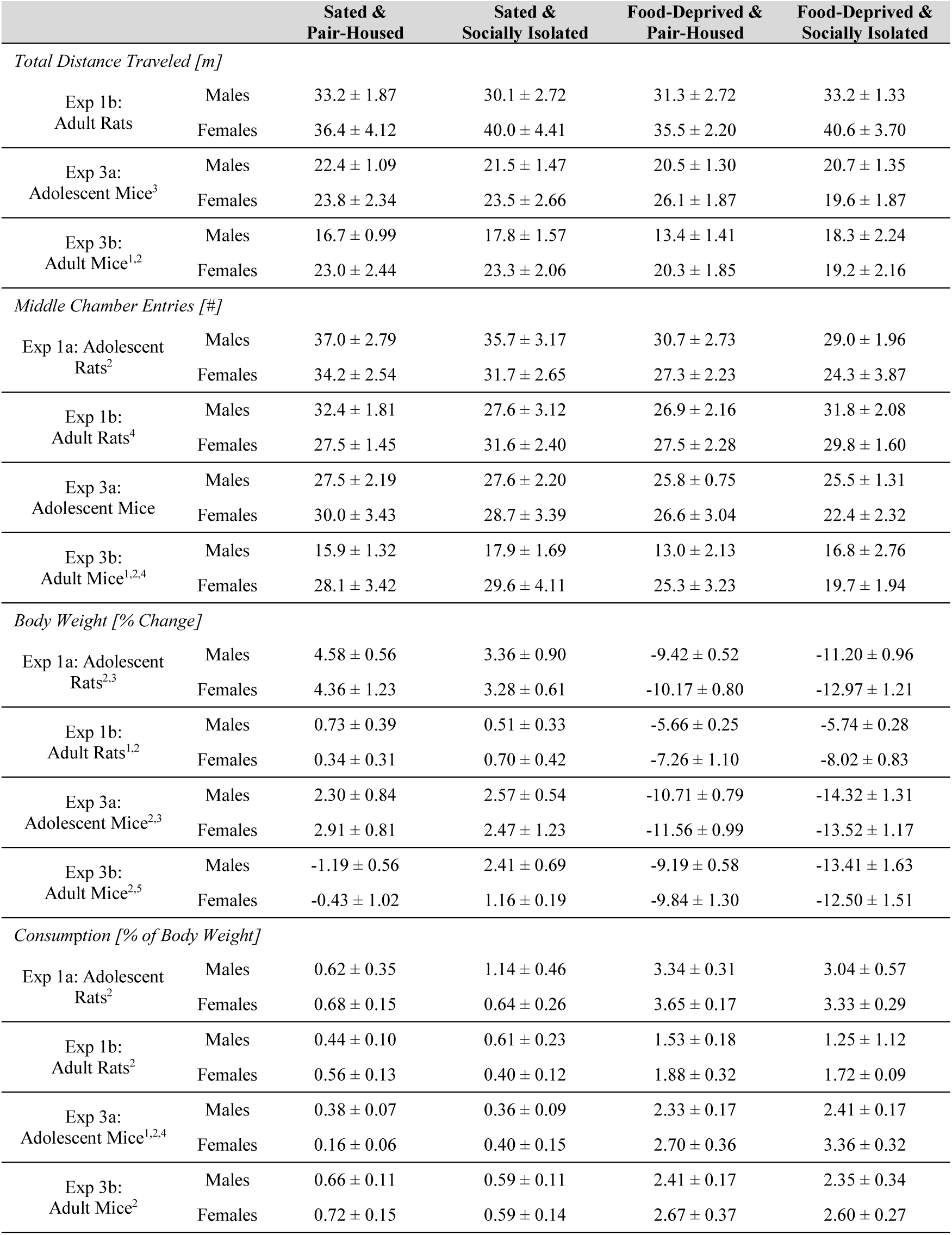
Activity measures, body weight, and post-test food consumption results for Experiments 1 and 3. Data shown as mean ± SEM. Distance traveled was not assessed in Experiment 1a due to technical difficulties. ^1^main effect of sex, ^2^main effect of hunger condition, ^3^main effect of housing condition, ^4^significant sex x hunger interaction, ^5^significant hunger x housing interaction, ^6^significant sex x hunger x housing interaction; see text for details and Tables 1, 3, 6, and 7 for corresponding ANOVA statistics.

|  |  | Sated &<br>Pair-Housed | Sated &<br>Socially Isolated | Food-Deprived &<br>Pair-Housed | Food-Deprived &<br>Socially Isolated |
| --- | --- | --- | --- | --- | --- |
| *Total Distance Traveled [m]* |  |  |  |  |  |
| Exp 1b:<br>Adult Rats | Males | 33.2 ± 1.87 | 30.1 ± 2.72 | 31.3 ± 2.72 | 33.2 ± 1.33 |
| | Females | 36.4 ± 4.12 | 40.0 ± 4.41 | 35.5 ± 2.20 | 40.6 ± 3.70 |
| Exp 3a:<br>Adolescent Mice<sup>3</sup> | Males | 22.4 ± 1.09 | 21.5 ± 1.47 | 20.5 ± 1.30 | 20.7 ± 1.35 |
| | Females | 23.8 ± 2.34 | 23.5 ± 2.66 | 26.1 ± 1.87 | 19.6 ± 1.87 |
| Exp 3b:<br>Adult Mice<sup>1,2</sup> | Males | 16.7 ± 0.99 | 17.8 ± 1.57 | 13.4 ± 1.41 | 18.3 ± 2.24 |
| | Females | 23.0 ± 2.44 | 23.3 ± 2.06 | 20.3 ± 1.85 | 19.2 ± 2.16 |
| *Middle Chamber Entries [#]* |  |  |  |  |  |
| Exp 1a: Adolescent<br>Rats<sup>2</sup> | Males | 37.0 ± 2.79 | 35.7 ± 3.17 | 30.7 ± 2.73 | 29.0 ± 1.96 |
| | Females | 34.2 ± 2.54 | 31.7 ± 2.65 | 27.3 ± 2.23 | 24.3 ± 3.87 |
| Exp 1b:<br>Adult Rats<sup>4</sup> | Males | 32.4 ± 1.81 | 27.6 ± 3.12 | 26.9 ± 2.16 | 31.8 ± 2.08 |
| | Females | 27.5 ± 1.45 | 31.6 ± 2.40 | 27.5 ± 2.28 | 29.8 ± 1.60 |
| Exp 3a:<br>Adolescent Mice | Males | 27.5 ± 2.19 | 27.6 ± 2.20 | 25.8 ± 0.75 | 25.5 ± 1.31 |
| | Females | 30.0 ± 3.43 | 28.7 ± 3.39 | 26.6 ± 3.04 | 22.4 ± 2.32 |
| Exp 3b:<br>Adult Mice<sup>1,2,4</sup> | Males | 15.9 ± 1.32 | 17.9 ± 1.69 | 13.0 ± 2.13 | 16.8 ± 2.76 |
| | Females | 28.1 ± 3.42 | 29.6 ± 4.11 | 25.3 ± 3.23 | 19.7 ± 1.94 |
| *Body Weight [% Change]* |  |  |  |  |  |
| Exp 1a: Adolescent<br>Rats<sup>2,3</sup> | Males | 4.58 ± 0.56 | 3.36 ± 0.90 | -9.42 ± 0.52 | -11.20 ± 0.96 |
| | Females | 4.36 ± 1.23 | 3.28 ± 0.61 | -10.17 ± 0.80 | -12.97 ± 1.21 |
| Exp 1b:<br>Adult Rats<sup>1,2</sup> | Males | 0.73 ± 0.39 | 0.51 ± 0.33 | -5.66 ± 0.25 | -5.74 ± 0.28 |
| | Females | 0.34 ± 0.31 | 0.70 ± 0.42 | -7.26 ± 1.10 | -8.02 ± 0.83 |
| Exp 3a:<br>Adolescent Mice<sup>2,3</sup> | Males | 2.30 ± 0.84 | 2.57 ± 0.54 | -10.71 ± 0.79 | -14.32 ± 1.31 |
| | Females | 2.91 ± 0.81 | 2.47 ± 1.23 | -11.56 ± 0.99 | -13.52 ± 1.17 |
| Exp 3b:<br>Adult Mice<sup>2,5</sup> | Males | -1.19 ± 0.56 | 2.41 ± 0.69 | -9.19 ± 0.58 | -13.41 ± 1.63 |
| | Females | -0.43 ± 1.02 | 1.16 ± 0.19 | -9.84 ± 1.30 | -12.50 ± 1.51 |
| *Consumption [% of Body Weight]* |  |  |  |  |  |
| Exp 1a: Adolescent<br>Rats<sup>2</sup> | Males | 0.62 ± 0.35 | 1.14 ± 0.46 | 3.34 ± 0.31 | 3.04 ± 0.57 |
| | Females | 0.68 ± 0.15 | 0.64 ± 0.26 | 3.65 ± 0.17 | 3.33 ± 0.29 |
| Exp 1b:<br>Adult Rats<sup>2</sup> | Males | 0.44 ± 0.10 | 0.61 ± 0.23 | 1.53 ± 0.18 | 1.25 ± 1.12 |
| | Females | 0.56 ± 0.13 | 0.40 ± 0.12 | 1.88 ± 0.32 | 1.72 ± 0.09 |
| Exp 3a:<br>Adolescent Mice<sup>1,2,4</sup> | Males | 0.38 ± 0.07 | 0.36 ± 0.09 | 2.33 ± 0.17 | 2.41 ± 0.17 |
| | Females | 0.16 ± 0.06 | 0.40 ± 0.15 | 2.70 ± 0.36 | 3.36 ± 0.32 |
| Exp 3b:<br>Adult Mice<sup>2</sup> | Males | 0.66 ± 0.11 | 0.59 ± 0.11 | 2.41 ± 0.17 | 2.35 ± 0.34 |
| | Females | 0.72 ± 0.15 | 0.59 ± 0.14 | 2.67 ± 0.37 | 2.60 ± 0.27 |

Adolescent rats lost weight under food deprivation and gained weight under sated conditions, resulting in a significant main effect of hunger condition on percent change in body weight (**Tables 1, 2**). There was also a significant main effect of housing condition on percent change in body weight; under socially isolated conditions adolescent rats lost more weight when food-deprived and gained less weight when sated compared to under pair-housed conditions (**Tables 1, 2**). Adolescent rats consumed significantly more food when food-deprived compared to when sated under both pair-housed and socially isolated conditions (**Tables 1, 2)**. There was no effect of sex on body weight or food consumption measures (**Tables 1, 2**).

### 3.2 Experiment 1b: Social versus food preference was altered by food deprivation, but not social isolation in adult rats

Adult rats had a significant reduction in social over food preference scores when tested under food-deprived compared to sated conditions; neither housing condition nor sex altered stimulus preference (**Table 3, Fig 3A, B**). Even though social over food preference was reduced following food deprivation, adult rats had a strong preference for the social stimulus under both sated (pair-housed: *t*_(15)_=10.72, *p* <0.001, *d* = 2.68; socially isolated: *t*_(15)_ = 10.10, *p* < 0.001, *d* = 2.52), and food-deprived (pair-housed: *t*_(15)_ = 3.81, *p* = 0.002, *d* = 0.95; socially isolated: *t*_(15)_ = 4.35, *p* = 0.001, *d* =1.09; **Fig 3A, B**) conditions.

**Table 3.** ANOVA statistics and partial eta squared (η^2^) effect sizes for Experiment 1b: Social versus food preference was altered by food deprivation, but not social isolation in adult rats. Significant effects shown in **bold**, n.s. = none significant, n.a. = not applicable, BW = body weight.

|  | Sex | Hunger Condition | Housing Condition | Interactions |
| --- | --- | --- | --- | --- |
| Social over Food Preference [%] | $F_{(1,14)} = 0.12, p = 0.73, \eta^2 = 0.009$ | $F_{(1,14)} = 8.83, p = 0.010, \eta^2 = 0.39$ | $F_{(1,14)} = 0.08, p = 0.78, \eta^2 = 0.006$ | n.s. |
| Social Stimulus Investigation [sec] | $F_{(1,14)} = 2.27, p = 0.15, \eta^2 = 0.14$ | $F_{(1,14)} = 2.48, p = 0.14, \eta^2 = 0.15$ | $F_{(1,14)} = 0.00, p = 0.99, \eta^2 = 0.99$ | Hunger x Housing:<br>$F_{(1,14)} = 5.62, p = 0.033, \eta^2 = 0.29$ |
| Food Stimulus Investigation [sec] | $F_{(1,14)} = 1.24, p = 0.28, \eta^2 = 0.082$ | $F_{(1,14)} = 16.1, p = 0.001, \eta^2 = 0.53$ | $F_{(1,14)} = 0.02, p = 0.89, \eta^2 = 0.001$ | n.s. |
| Total Investigation [sec] | $F_{(1,14)} = 3.57, p = 0.080, \eta^2 = 0.20$ | $F_{(1,14)} = 21.8, p < 0.001, \eta^2 = 0.61$ | $F_{(1,14)} = 0.005, p = 0.95, \eta^2 < 0.001$ | Hunger x Housing:<br>$F_{(1,14)} = 5.21, p = 0.039, \eta^2 = 0.27$ |
| Total Distance [m] | $F_{(1,9)} = 4.22, p = 0.070, \eta^2 = 0.32$ | $F_{(1,9)} = 0.03, p = 0.87, \eta^2 = 0.003$ | $F_{(1,9)} = 1.36, p = 0.27, \eta^2 = 0.13$ | n.s. |
| Middle Chamber Entries [#] | $F_{(1,14)} = 0.06, p = 0.81, \eta^2 = 0.004$ | $F_{(1,14)} = 0.47, p = 0.50, \eta^2 = 0.033$ | $F_{(1,14)} = 2.06, p = 0.17, \eta^2 = 0.13$ | Sex x Hunger x Housing:<br>$F_{(1,14)} = 5.34, p = 0.037, \eta^2 = 0.28$ |
| Body Weight [% change] | $F_{(1,14)} = 8.89, p = 0.010, \eta^2 = 0.39$ | $F_{(1,14)} = 298.7, p < 0.001, \eta^2 = 0.96$ | $F_{(1,14)} = 0.24, p = 0.63, \eta^2 = 0.017$ | n.s. |
| Consumption [% of BW] – Pair-Housed | $F_{(1,6)} = 0.94, p = 0.37, \eta^2 = 0.14$ | $F_{(1,6)} = 67.5, p < 0.001, \eta^2 = 0.92$ | n.a. | n.s. |
| Consumption [% of BW] – Socially Isolated | $F_{(1,14)} = 1.14, p = 0.30, \eta^2 = 0.075$ | $F_{(1,14)} = 30.9, p < 0.001, \eta^2 = 0.69$ | n.a. | n.s. |

**Fig 3.**
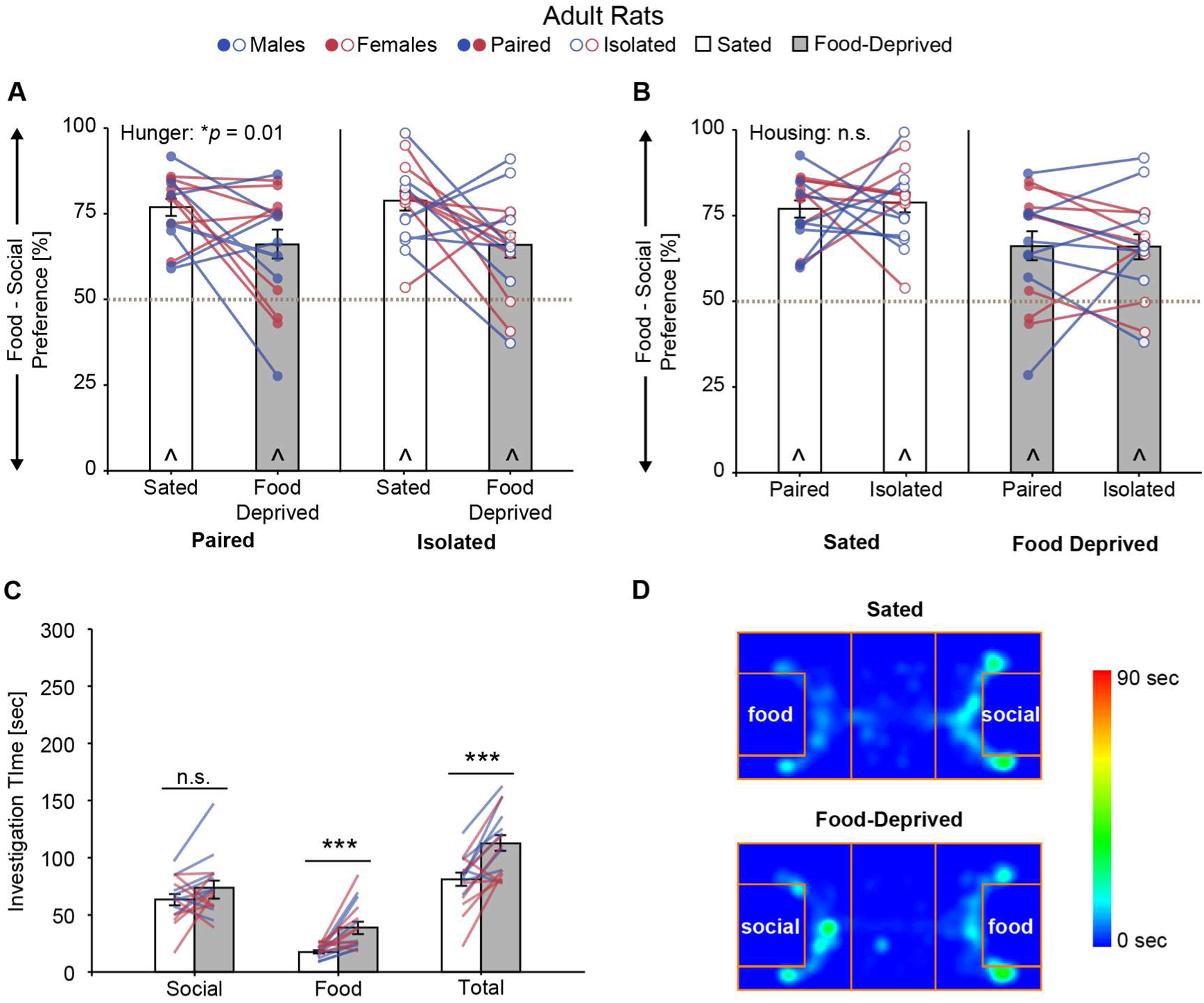
Experiment 1b. Food deprivation (**A**), but not social isolation (**B**), significantly attenuated preference for the social stimulus in adult rats. Food deprivation did not alter investigation of the social stimulus, but robustly increased investigation of the food stimulus (**C**). Representative heat maps of activity from one subject (**D**). Bar graphs (mean ± SEM) collapsed across sex for **A** and **B** (same data replotted to illustrate main effects), and across sex and housing condition for **C**; individual data collapsed across housing condition in **C**; * *p* < 0.05, ** *p* < 0.01, *** *p* < 0.001 mixed-model ANOVA; ^ p < 0.05, one-sample t-Test from 50% (gray dashed line); n.s. = not significant; n = 8 males, n = 8 females.

When food-deprived, adult rats increased their investigation of the food stimulus resulting in an increase in total investigation time compared to when they were sated (**Fig 3C, Table 3**). There were no main effects of housing condition nor sex on stimulus investigation times in adult rats, however there were significant hunger condition by housing condition interactions on social and total investigation times (**Fig 3C, Table 3**). *Post hoc* comparisons showed that for both of these measures adult rats spent more time investigating under food-deprived compared to sated conditions when they had been socially isolated (social: sated = 58.1 ± 4.60 sec, food-deprived = 79.5 ± 7.27 sec, *p* = 0.028, total: sated = 73.3 ± 5.14 sec, food-deprived = 121.69 ± 9.53 sec, *p* = 0.001), but not when they had been pair-housed (social: sated = 68.3 ± 8.49 sec, food-deprived = 69.1± 7.70 sec, *p* = 0.91, total: sated = 89.44 ± 8.70 sec, food-deprived = 104 ± 10.1 sec, *p* = 0.09); no other paired comparisons reached significance (*p* > 0.05, all).

Locomotor activity, as measured by distance traveled, was similar across all conditions and between both sexes in adult rats (**Tables 2, 3**). However, there was a significant sex by hunger condition by housing condition interaction on locomotor activity as measured by middle chamber entries (**Tables 2, 3**). *Post hoc* comparisons showed that under food-deprived conditions adult males had fewer middle chamber entries when they had been pair-housed compared to when they had been socially isolated (*p* = 0.04), and that under pair-housed conditions adult males had more middle chamber entries when they were sated compared to when they were food-deprived (*p* = 0.025); no other paired comparisons reached significance (*p* > 0.05, all).

There was a significant main effect of hunger condition on percent change in body weight with adult rats losing weight under food-deprived but not under sated conditions (**Tables 2, 3**). There were no effects of housing condition on percent change in body weight, but the percent change in body weight was greater in adult females than adult males (**Tables 2, 3**). Under both pair-housed and socially isolated conditions, adult rats consumed significantly more food when food-deprived compared to when sated (**Tables 2, 3**). There was no effect of sex on food consumption (**Tables 2, 3**).

### 3.3 Experiment 2: Time-of-testing did not alter food deprivation-driven changes in social versus food preference in adolescent rats

Adolescent rats had significantly lower social over food preference scores under food-deprived compared to under sated conditions, while time-of-testing did not affect stimulus preference (**Table 4, Fig 4A, B**). There was a significant sex by hunger condition interaction on stimulus preference (**Table 4, Fig 4A**). *Post hoc* comparisons showed that under sated conditions stimulus preference was similar between males and females (*p* = 0.31), but that under food-deprived conditions social over food preference was lower in adolescent females than adolescent males (*p* = 0.032). When collapsed acorss sex, adolescent rats showed a significant social preference under sated conditions (dark phase: *t*_(13)_ = 38.8, *p* < 0.001, *d* = 8.59; light phase: *t*_(13)_ = 27.4, *p* < 0.001, *d* = 6.66) and no stimulus preference under food-deprived conditions (dark phase: *t*_(13)_ = 1.53, *p* = 0.15, *d* = 0.021; light phase: *t*_(13)_ = 1.44, *p* = 0.17, *d* = 0.064; **Fig 4A, B**).

**Table 4.** ANOVA statistics and partial eta squared (η^2^) effect sizes for Experiment 2: Time-of-testing did not alter food deprivation-driven changes in social versus food preference in adolescent rats. Significant effects shown in **bold**, n.s. = none significant, BW = body weight.

|  | Sex | Hunger Condition | Time-of-testing | Interactions |
| --- | --- | --- | --- | --- |
| Social over Food Preference [%] | $F_{(1,12)} = 5.93, p = 0.031, \eta^2 = 0.33$ | $F_{(1,12)} = 82.1, p < 0.001, \eta^2 = 0.87$ | $F_{(1,12)} = 0.38, p = 0.55, \eta^2 = 0.031$ | Sex x Hunger:<br>$F_{(1,12)} = 4.81, p = 0.049, \eta^2 = 0.29$ |
| Social Stimulus Investigation [sec] | $F_{(1,12)} = 25.1, p < 0.001, \eta^2 = 0.67$ | $F_{(1,12)} = 14.9, p = 0.002, \eta^2 = 0.55$ | $F_{(1,12)} = 1.57, p = 0.24, \eta^2 = 0.12$ | n.s. |
| Food Stimulus Investigation [sec] | $F_{(1,12)} = 0.76, p = 0.40, \eta^2 = 0.059$ | $F_{(1,12)} = 56.9, p < 0.001, \eta^2 = 0.83$ | $F_{(1,12)} = 2.02, p = 0.18, \eta^2 = 0.14$ | n.s. |
| Total Investigation [sec] | $F_{(1,12)} = 10.1, p = 0.008, \eta^2 = 0.46$ | $F_{(1,12)} = 7.08, p = 0.021, \eta^2 = 0.37$ | $F_{(1,12)} = 3.41, p = 0.090, \eta^2 = 0.22$ | n.s. |
| Total Distance [m] | $F_{(1,10)} = 0.70, p = 0.42, \eta^2 = 0.066$ | $F_{(1,10)} = 0.73, p = 0.41, \eta^2 = 0.068$ | $F_{(1,10)} = 3.92, p = 0.076, \eta^2 = 0.28$ | n.s. |
| Middle Chamber Entries [#] | $F_{(1,12)} = 1.75, p = 0.21, \eta^2 = 0.13$ | $F_{(1,12)} = 6.99, p = 0.021, \eta^2 = 0.37$ | $F_{(1,12)} = 0.45, p = 0.52, \eta^2 = 0.036$ | n.s. |
| Body Weight [% change] | $F_{(1,12)} = 13.8, p = 0.003, \eta^2 = 0.53$ | $F_{(1,12)} = 488, p < 0.001, \eta^2 = 0.98$ | $F_{(1,12)} = 3.47, p = 0.087, \eta^2 = 0.22$ | Sex x Hunger:<br>$F_{(1,12)} = 8.74, p = 0.012, \eta^2 = 0.42$ |
| Consumption [% of BW] | $F_{(1,12)} = 2.28, p = 0.16, \eta^2 = 0.16$ | $F_{(1,12)} = 165, p < 0.001, \eta^2 = 0.93$ | $F_{(1,12)} = 0.29, p = 0.60, \eta^2 = 0.023$ | n.s. |

**Fig 4.**
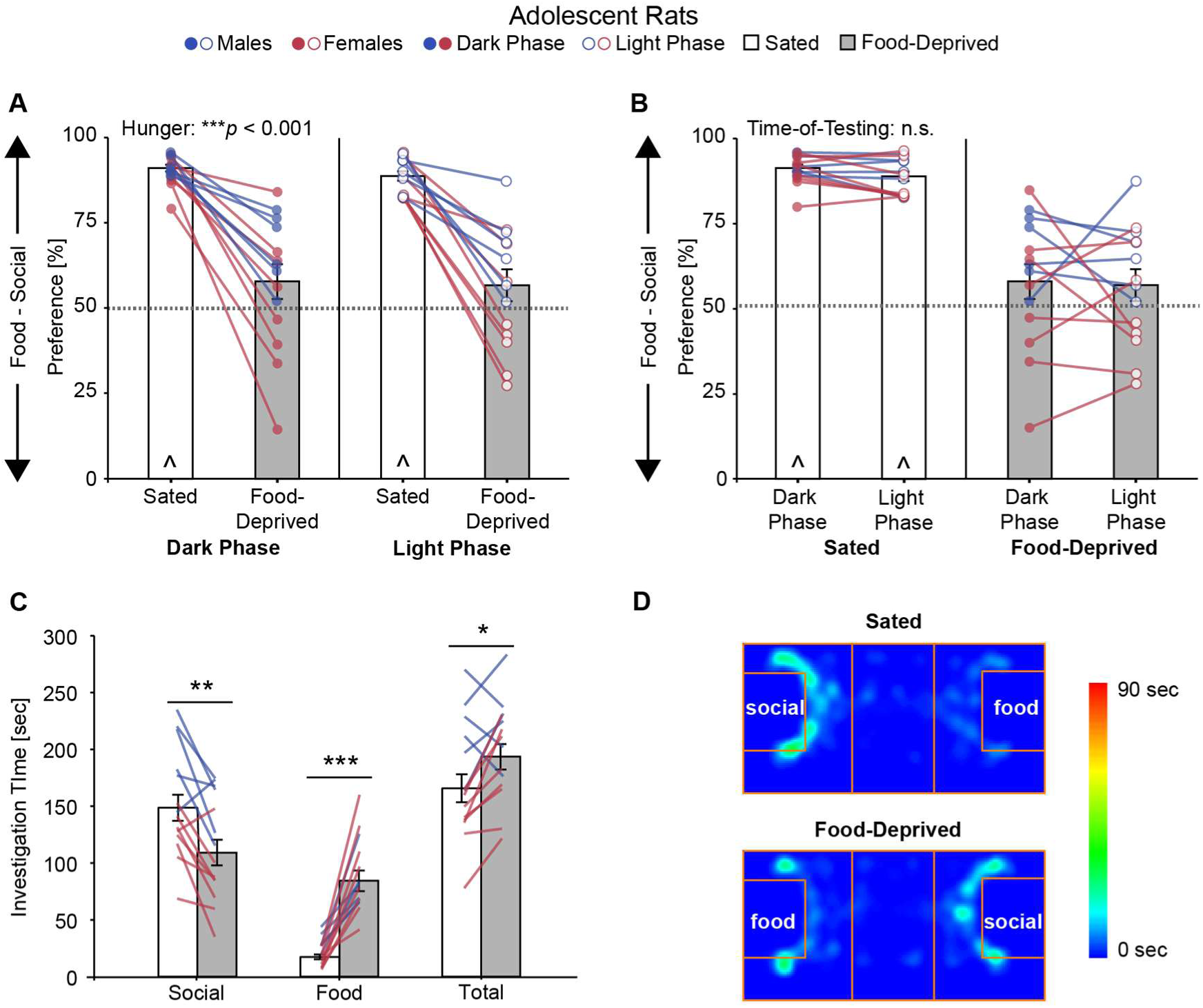
Experiment 2. Food deprivation abolished preference for the social stimulus in adolescent rats (**A**), while time-of-testing had no effect of stimulus preference (**B**). Food deprivation decreased investigation of the social stimulus, and robustly increased investigation of the food stimulus (**C**). Representative heat maps of activity from one subject (**D**). Bar graphs (mean ± SEM) collapsed across sex for **A** and **B** (same data replotted to illustrate main effects), and across sex and time-of-testing for **C**; individual data collapsed across time-of-testing condition in **C**; * *p* < 0.05, ** *p* < 0.01, *** *p* < 0.001 mixed-model ANOVA; ^ p < 0.05, one-sample t-Test from 50% (gray dashed line); n.s. = not significant; n = 6 males, n = 8 females.

The observed changes in stimulus preference were due to decreased time spent investigating the social stimulus and increased time spent investigating the food stimulus when adolescent rats were food-deprived compared to when they were sated (**Fig 4C, Table 4**). There was a mild, but significant, increase in total investigation time under food-deprived compared to sated conditions (**Fig 4C, Table 4**). There were no sex differences in time spent investigating the food stimulus, but adolescent males spent more time investigating the social stimulus than adolescent females resulting in a greater total investigation time in adolescent males compared to adolescent females (**Fig 4C, Table 4**). Stimulus investigation times were unaffected by time-of-testing in adolescent rats (**Table 4**).

Locomotor activity, as measured by middle chamber entries, was decreased under food-deprived compared to sated conditions and unaffected by time-of-testing, however distance traveled was similar across all conditions. Locomotor activity was similar between adolescent male and female rats for both measures (**Tables 4, 5**).

**Table 5.** Activity measures, body weight, and post-test food consumption results for Experiment 2. Data shown as mean ± SEM. ^1^main effect of sex, ^2^main effect of hunger condition, ^3^significant sex x hunger interaction; see text for details and Table 4 for corresponding ANOVA statistics.

|  | Sated & Dark Phase | Sated & Light Phase | Food-Deprived & Dark Phase | Food-Deprived & Light Phase |
| --- | --- | --- | --- | --- |
| *Total Distance Traveled [m]* |  |  |  |  |
| Males | 36.6 ± 2.35 | 28.8 ± 1.85 | 33.8 ± 1.77 | 31.6 ± 1.94 |
| Females | 38.0 ± 1.72 | 35.8 ± 4.88 | 34.0 ± 4.17 | 33.2 ± 3.02 |
| *Middle Chamber Entries [#]<sup>2</sup>* |  |  |  |  |
| Males | 27.0 ± 2.82 | 25.0 ± 2.46 | 24.7 ± 2.23 | 25.3 ± 3.03 |
| Females | 33.1 ± 1.33 | 31.0 ± 3.45 | 26.8 ± 3.23 | 24.9 ± 1.69 |
| *Body Weight [% Change]<sup>1,2,3</sup>* |  |  |  |  |
| Males | 3.44 ± 0.61 | 2.44 ± 0.49 | -7.66 ± 1.24 | -8.64 ± 0.91 |
| Females | 4.26 ± 0.74 | 2.67 ± 0.60 | -10.76 ± 0.57 | -11.34 ± 0.45 |
| *Consumption [% of Body Weight]<sup>2</sup>* |  |  |  |  |
| Males | 1.14 ± 0.34 | 1.10 ± 0.12 | 2.78 ± 0.26 | 3.22 ± 0.16 |
| Females | 0.67 ± 0.19 | 0.41 ± 0.10 | 3.09 ± 0.19 | 3.20 ± 0.20 |

Adolescent rats lost weight under food deprivation and gained weight under sated conditions resulting in a significant main effect of hunger condition on percent change in body weight (**Tables 4, 5**). There was a significant sex by hunger condition interaction on percent change in body weight (**Tables 4, 5**). *Post hoc* comparisons showed that body weight gain was similar between males and females under sated conditions (*p* = 0.49), but that body weight loss was greater in females than males under food-deprived conditions (*p* < 0.001). Time-of-testing did not affect percent change in body weight (**Tables 4, 5**). During both the light phase and the dark phase, adolescent rats consumed significantly more food when food-deprived compared to when sated (**Tables 4, 5**); there was no effect of time-of-testing or sex on food consumption (**Tables 4, 5**).

### 3.4 Experiment 3a: Social versus food preference was altered by food deprivation, but not social isolation in adolescent mice

Adolescent mice had a significant reduction in their social over food preference scores when tested under food-deprived compared to sated conditions; neither housing condition nor sex altered stimulus preference (**Table 6, Fig 5A, B**). Adolescent mice did not have a significant preference for either stimulus under sated conditions (pair-housed: *t*_(14)_= 0.45, *p* = 0.66, *d* = 0.12; isolated: *t*_(14)_ = 1.78, *p* = 0.097, *d* = 0.46), and food deprivation resulted in the emergence of a food preference (pair-housed: *t*_(14)_ = 15.18, *p* < 0.001, *d* = 3.92; isolated: *t*_(14)_ = 12.14, *p* < 0.001, *d* = 3.13; **Fig 5A, B**).

**Table 6.** ANOVA statistics and partial eta squared (η^2^) effect sizes for Experiment 3a: Social versus food preference was altered by food deprivation, but not social isolation in adolescent mice. Significant effects shown in **bold**, n.s. = none significant, BW = body weight.

|  | Sex | Hunger Condition | Housing Condition | Interactions |
| --- | --- | --- | --- | --- |
| Social over Food Preference [%] | $F_{(1,13)} = 0.37, p = 0.55, \eta^2 = 0.028$ | $F_{(1,13)} = \mathbf{140}, p < \mathbf{0.001}, \eta^2 = \mathbf{0.92}$ | $F_{(1,13)} = 0.52, p = 0.48, \eta^2 = 0.039$ | n.s. |
| Social Stimulus Investigation [sec] | $F_{(1,13)} = 4.10, p = 0.064, \eta^2 = 0.24$ | $F_{(1,13)} = \mathbf{7.90}, p = \mathbf{0.015}, \eta^2 = \mathbf{0.38}$ | $F_{(1,13)} = 0.76, p = 0.40, \eta^2 = 0.055$ | Sex x Hunger x Housing:<br>$F_{(1,13)} = \mathbf{5.45}, p = \mathbf{0.036}, \eta^2 = \mathbf{0.30}$ |
| Food Stimulus Investigation [sec] | $F_{(1,13)} = 1.20, p = 0.29, \eta^2 = 0.084$ | $F_{(1,13)} = \mathbf{94.59}, p < \mathbf{0.001}, \eta^2 = \mathbf{0.88}$ | $F_{(1,13)} = 3.34, p = 0.091, \eta^2 = 0.20$ | n.s. |
| Total Investigation [sec] | $F_{(1,13)} = 3.18, p = 0.10, \eta^2 = 0.20$ | $F_{(1,13)} = \mathbf{78.1}, p < \mathbf{0.001}, \eta^2 = \mathbf{0.86}$ | $F_{(1,13)} = 4.46, p = 0.055, \eta^2 = 0.26$ | n.s. |
| Total Distance [m] | $F_{(1,9)} = 1.40, p = 0.27, \eta^2 = 0.14$ | $F_{(1,9)} = 0.81, p = 0.39, \eta^2 = 0.082$ | $F_{(1,9)} = \mathbf{5.55}, p = \mathbf{0.043}, \eta^2 = \mathbf{0.38}$ | n.s. |
| Middle Chamber Entries [#] | $F_{(1,13)} = 0.006, p = 0.94, \eta^2 < 0.001$ | $F_{(1,13)} = 4.54, p = 0.053, \eta^2 = 0.26$ | $F_{(1,13)} = 1.34, p = 0.27, \eta^2 = 0.093$ | n.s. |
| Body Weight [% change] | $F_{(1,13)} = 0.015, p = 0.90, \eta^2 = 0.001$ | $F_{(1,13)} = \mathbf{1.13}, p < \mathbf{0.001}, \eta^2 = \mathbf{0.99}$ | $F_{(1,13)} = \mathbf{8.69}, p = \mathbf{0.011}, \eta^2 = \mathbf{0.40}$ | n.s. |
| Consumption [% of BW] | $F_{(1,13)} = \mathbf{15.1}, p = \mathbf{0.002}, \eta^2 = \mathbf{0.54}$ | $F_{(1,13)} = \mathbf{344.0}, p < \mathbf{0.001}, \eta^2 = \mathbf{0.96}$ | $F_{(1,13)} = 2.64, p = 0.13, \eta^2 = 1.17$ | Sex x Hunger<br>$F_{(1,13)} = \mathbf{8.63}, p = \mathbf{0.012}, \eta^2 = \mathbf{0.40}$ |

**Fig 5.**
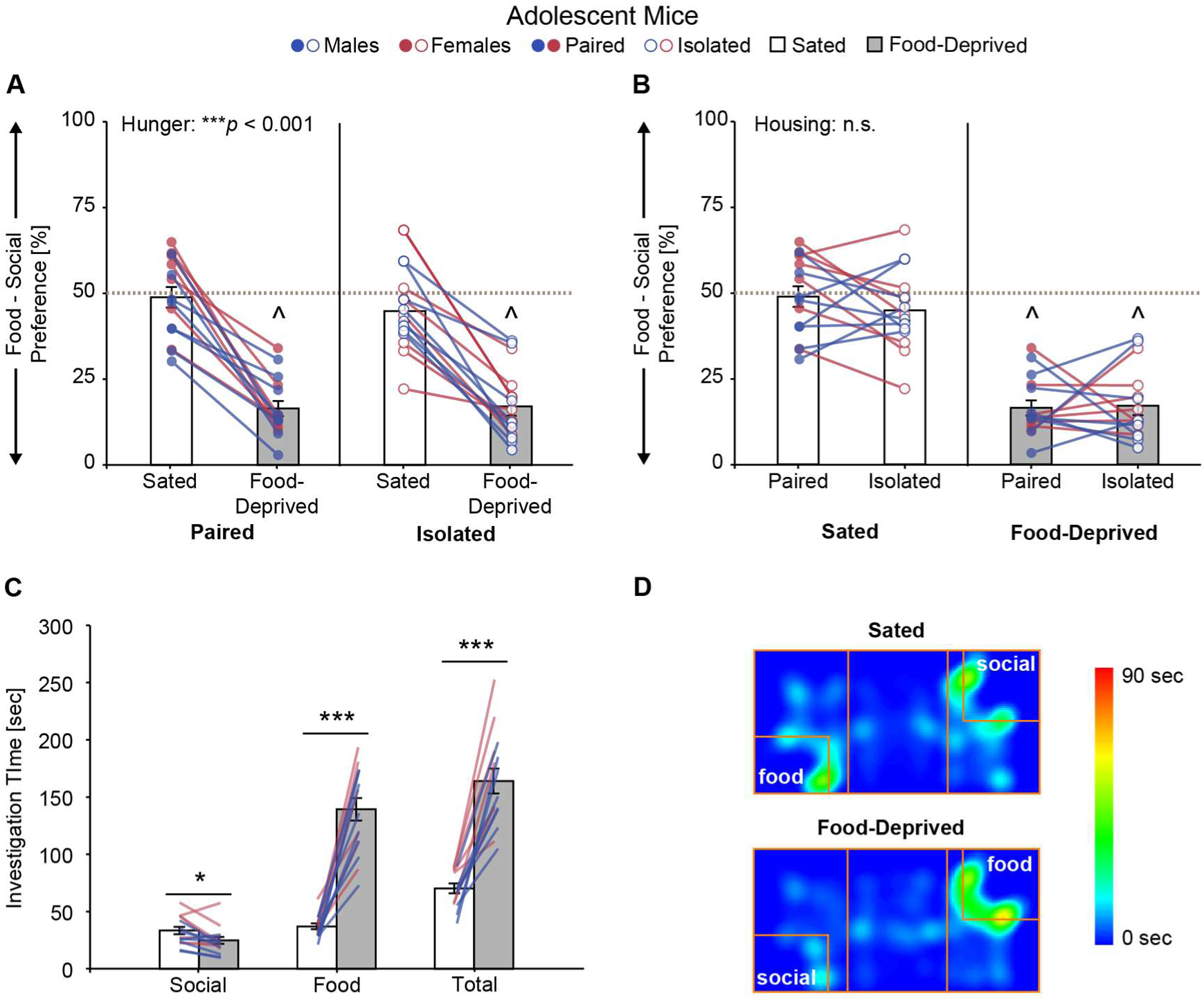
Experiment 3a. Food deprivation (**A**), but not social isolation (**B**), significantly altered stimulus preference in adolescent mice which resulted in the emergence of a food preference under food-deprived conditions. Food deprivation mildly decreased investigation of the social stimulus and robustly increased investigation of the food stimulus (**C**). Representative heat maps of activity from one subject (**D**). Bar graphs (mean ± SEM) collapsed across sex for **A** and **B** (same data replotted to illustrate main effects), and across sex and housing condition for **C**; individual data collapsed across housing condition in **C**; * *p* < 0.05, ** *p* < 0.01, *** *p* < 0.001 mixed-model ANOVA; ^ p < 0.05, one-sample t-Test from 50% (gray dashed line); n.s. = not significant; n = 8 males, n = 7 females.

The observed changes in stimulus preference were due to a mild, but significant, decrease in the time spent investigating the social stimulus and a robust increase in the time spent investigating the food stimulus which increased total investigation time when adolescent mice were food-deprived compared to when they were sated (**Table 6, Fig 5**). Food and total investigation times were unaffected by housing condition or sex in adolescent mice, but there was a significant sex by hunger condition by housing condition interaction on social investigation times (**Table 6**). *Post hoc* comparisons showed that pair-housed adolescent female mice investigated the social stimulus less when they were food-deprived compared to when they were sated (food-deprived: 26.1 ± 4.69, sated: 42.5 ± 4.13, *p* = 0.011), and that when subjects were tested under pair-housed and sated conditions adolescent male mice investigated the social stimulus less than adolescent female mice (males: 25.1 ± 3.87 sec, females: 42.5 ± 4.13 sec, *p* = 0.009); no other paired comparisons reached significance (*p* > 0.05, all).

Adolescent mice had a tendency to make more middle chamber entries when food-deprived compared to when sated, but distance traveled was similar across hunger conditions (**Tables 2, 6**). Adolescent mice traveled further under pair-housed compared to isolated conditions, but middle chamber entries were similar across housing conditions (**Tables 2, 6**). There was no effect of sex on either locomotor activity measure in adolescent mice (**Tables 2, 6**).

As expected, adolescent mice lost weight under food-deprived conditions and gained weight under sated conditions resulting in a significant main effect of hunger condition on percent change in body weight (**Tables 2, 6**). Interestingly, there was also a significant main effect of housing condition on percent change in body weight; adolescent mice lost more weight when food-deprived and gained less weight when sated when tested under socially isolated conditions compared to under pair-housed conditions (**Tables 2, 6**). There was no effect of sex on percent change in body weight in adolescent mice (**Tables 2, 6**). As expected, adolescent mice consumed significantly more food when food-deprived compared to when sated (**Tables 2, 6**). There was a significant sex by hunger condition interaction on food consumption (**Tables 2, 6**). *Post hoc* comparisons showed that adolescent female mice consumed more food relative to their body weight than adolescent male mice when subjects were food-deprived (*p* = 0.002); the sexes consumed a similar amount of food when subjects were sated (*p* = 0.43). There was no effect of housing condition on food consumption in adolescent mice (**Tables 2, 6**).

### 3.5 Experiment 3b: Social versus food preference was altered by food deprivation, but not social isolation in adult mice

Adult mice had a significant reduction in their social over food preference scores when tested under food-deprived compared to sated conditions; neither housing condition nor sex altered stimulus preference (**Table 7, Fig 6A, B**). Adult mice did not have a significant preference for either stimulus under sated conditions (pair-housed: *t*_(14)_= 0.61, *p* = 0.55, *d* = 0.16; isolated: *t*_(14)_ = 1.44, *p* = 0.17, *d* = 0.37) while food deprivation resulted in the emergence of a food preference (pair-housed: *t*_(14)_ = 9.09, *p* < 0.001, *d* = 2.35; isolated: *t*_(14)_ = 5.26, *p* < 0.001, *d* = 1.36; **Fig 6A, B**).

**Table 7.** ANOVA statistics and partial eta squared (η^2^) effect sizes for Experiment 3b: Social versus food preference was altered by food deprivation, but not social isolation in adult mice. Significant effects shown in **bold**, n.s. = none significant, BW = body weight.

|  | Sex | Hunger Condition | Housing Condition | Interactions |
| --- | --- | --- | --- | --- |
| Social over Food Preference [%] | $F_{(1,13)} = 1.94, p = 0.19, \eta^2 = 0.13$ | $F_{(1,13)} = \mathbf{44.8}, p < \mathbf{0.001}, \eta^2 = \mathbf{0.78}$ | $F_{(1,13)} = 0.014, p = 0.91, \eta^2 = 0.001$ | n.s. |
| Social Stimulus Investigation [sec] | $F_{(1,13)} = 1.86, p = 0.20, \eta^2 = 0.13$ | $F_{(1,13)} = 0.67, p = 0.43, \eta^2 = 0.049$ | $F_{(1,13)} = 0.37, p = 0.56, \eta^2 = 0.027$ | n.s. |
| Food Stimulus Investigation [sec] | $F_{(1,13)} = 0.008, p = 0.93, \eta^2 = 0.001$ | $F_{(1,13)} = \mathbf{41.1}, p < \mathbf{0.001}, \eta^2 = \mathbf{0.76}$ | $F_{(1,13)} = 0.47, p = 0.51, \eta^2 = 0.035$ | n.s. |
| Total Investigation [sec] | $F_{(1,13)} = 0.18, p = 0.68, \eta^2 = 0.014$ | $F_{(1,13)} = \mathbf{43.5}, p < \mathbf{0.001}, \eta^2 = \mathbf{0.77}$ | $F_{(1,13)} = 1.43, p = 0.25, \eta^2 = 0.099$ | n.s. |
| Total Distance [m] | $F_{(1,12)} = \mathbf{5.98}, p = \mathbf{0.03}, \eta^2 = \mathbf{0.33}$ | $F_{(1,12)} = \mathbf{7.90}, p = \mathbf{0.016}, \eta^2 = \mathbf{0.40}$ | $F_{(1,12)} = 1.32, p = 0.27, \eta^2 = 0.099$ | n.s. |
| Middle Chamber Entries [#] | $F_{(1,13)} = \mathbf{12.89}, p = \mathbf{0.003}, \eta^2 = \mathbf{0.50}$ | $F_{(1,13)} = \mathbf{17.4}, p = \mathbf{0.001}, \eta^2 = \mathbf{0.57}$ | $F_{(1,13)} = 0.06, p = 0.82, \eta^2 = 0.004$ | Hunger x Sex:<br>$F_{(1,13)} = \mathbf{4.72}, p = \mathbf{0.049}, \eta^2 = \mathbf{0.27}$ |
| Body Weight [% change] | $F_{(1,13)} = 0.004, p = 0.95, \eta^2 < 0.001$ | $F_{(1,13)} = \mathbf{272}, p < \mathbf{0.001}, \eta^2 = \mathbf{0.95}$ | $F_{(1,13)} = 0.36, p = 0.56, \eta^2 = 0.027$ | Hunger x Housing:<br>$F_{(1,13)} = \mathbf{18.4}, p = \mathbf{0.001}, \eta^2 = \mathbf{0.59}$ |
| Consumption [% of BW] | $F_{(1,13)} = 1.01, p = 0.33, \eta^2 = 0.072$ | $F_{(1,13)} = \mathbf{147}, p < \mathbf{0.001}, \eta^2 = \mathbf{0.92}$ | $F_{(1,13)} = 0.31, p = 0.59, \eta^2 = 0.023$ | n.s. |

**Fig 6.**
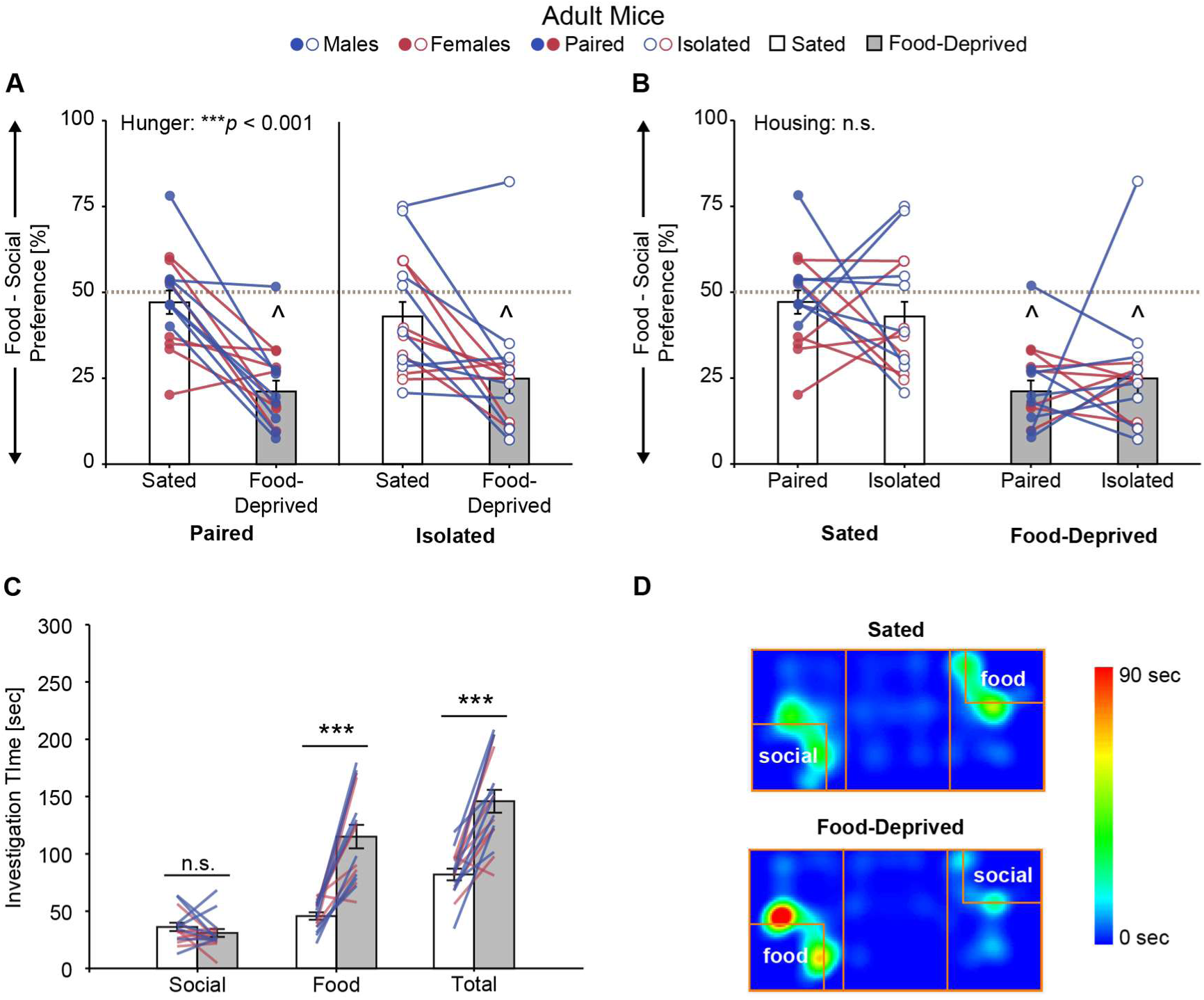
Experiment 3b. Food deprivation (**A**), but not social isolation (**B**), significantly altered stimulus preference in adult mice, which resulted in the emergence of a food preference under food-deprived conditions. Food deprivation robustly increased investigation of the food stimulus, but did not change investigation of the social stimulus (**C**). Representative heat maps of activity from one subject (**D**). Bar graphs (mean ± SEM) collapsed across sex for **A** and **B** (same data replotted to illustrate main effects), and across sex and housing condition for **C**; individual data collapsed across housing condition in **C**; *** *p* < 0.001 mixed-model ANOVA; ^ p < 0.05, one-sample t-Test from 50% (gray dashed line); n.s. = not significant; n = 8 males, n = 7 females.

The observed changes in stimulus preference were due to a robust increase in the time spent investigating the food stimulus, which increased total investigation time when adult mice were food-deprived compared to when they were sated; time spent investigating the social stimulus was similar across conditions (**Table 7, Fig 5**). Investigation times were unaffected by housing condition or sex in adult mice (**Table 7**).

There was a main effect of sex on locomotor activity as measured by distance traveled; adult female mice traveled further than adult male mice (**Tables 2, 7**). There was also a main effect of hunger condition on distance traveled; adult mice traveled further when sated compared to when they were food-deprived (**Tables 2, 7**). There was a significant sex by hunger condition interaction on locomotor activity as measured by middle chamber entries, but not by distance traveled. *Post hoc* paired comparisons showed that adult female mice (*p* = 0.001), but not adult male mice (*p* = 0.17), made more chamber entries when sated compared to when food-deprived. Further, similar to distance traveled, adult female mice made more middle entries than adult male mice when sated (*p* = 0.001) and when food-deprived (*p* = 0.024). Locomotor activity measures were unaffected by housing condition in adult mice (**Tables 2, 7**).

As expected, there was a significant main effect of hunger condition on percent change in body weight; adult mice lost weight under food-deprived compared to under sated conditions (**Tables 2, 7**). There was also a significant hunger condition by housing condition interaction on percent change in body weight (**Tables 2, 7**). *Post hoc* comparisons showed that when adult mice were socially isolated compared to when they were pair-housed, they lost more weight when food-deprived (*p* = 0.018), and gained more when they were sated (*p* = 0.001). These patterns were similar between males and females; there was no effect or interaction with sex on percent change in body weight (**Tables 2, 7**). Adult mice consumed significantly more food relative to their body weight when food-deprived compared to when sated; there were no effects of sex or housing condition on food consumption (**Tables 2, 7**).

### 3.6 Experiment 4a: Social over food preference was greater for a novel social stimulus compared to a familiar social stimulus in adolescent rats

Adolescent rats had higher social over food preference scores when the social stimulus was novel compared to when the social stimulus was their cagemate (**Fig 7A, Table 8)**; there was no effect of sex on stimulus preference. Adolescent rats exhibited a robust social preference, and this was true for both the novel (*t*_(16)_ = 14.28, *p* < 0.001, *d* = 3.46) and familiar (cagemate; *t*_(16)_ = 7.04, *p* < 0.001, *d* = 1.71) social stimuli (**Fig 7A**).

**Table 8.**
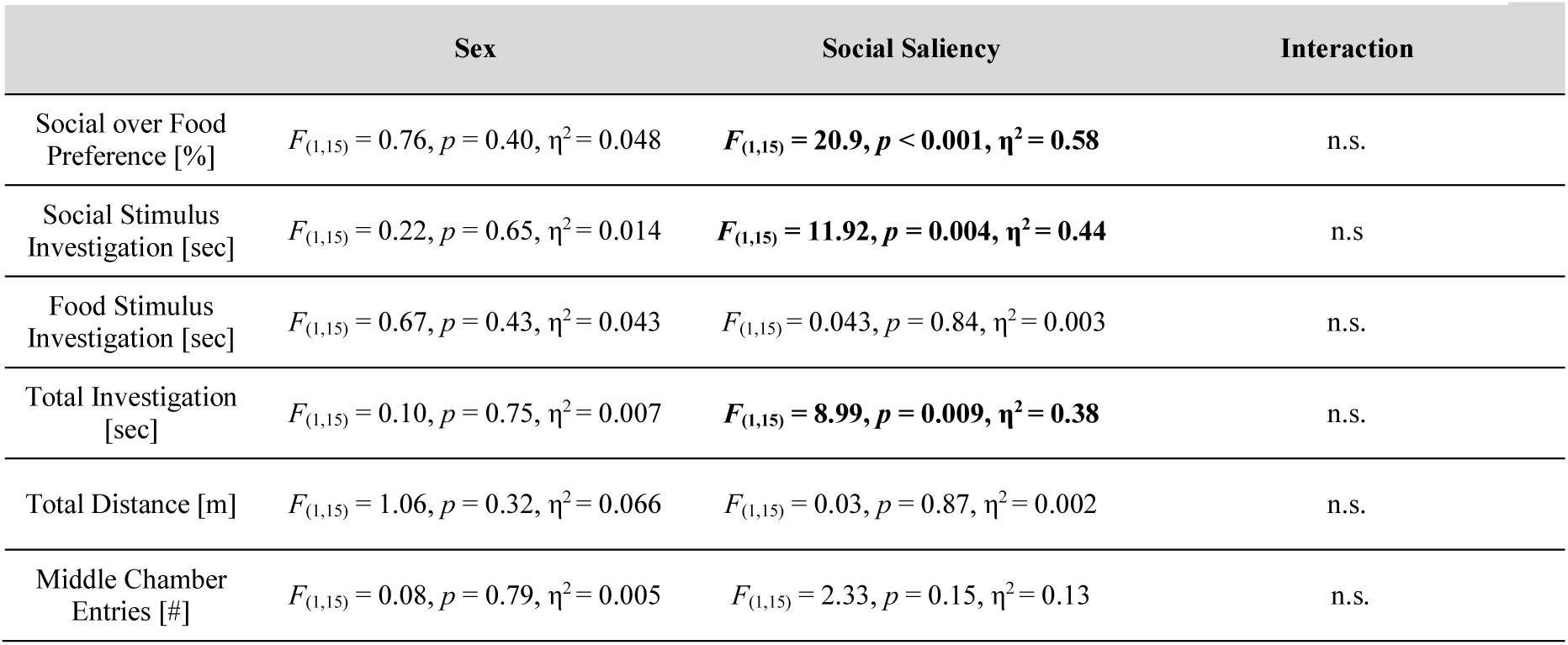
ANOVA statistics and partial eta squared (η^2^) effect sizes for Experiment 4a: Social over food preference was greater for a novel social stimulus compared to a familiar social stimulus in adolescent rats. Significant effects shown in **bold**, n.s. = none significant.

|  | Sex | Social Saliency | Interaction |
| --- | --- | --- | --- |
| Social over Food Preference [%] | $F_{(1,15)} = 0.76, p = 0.40, \eta^2 = 0.048$ | $F_{(1,15)} = \mathbf{20.9}, p < \mathbf{0.001}, \eta^2 = \mathbf{0.58}$ | n.s. |
| Social Stimulus Investigation [sec] | $F_{(1,15)} = 0.22, p = 0.65, \eta^2 = 0.014$ | $F_{(1,15)} = \mathbf{11.92}, p = \mathbf{0.004}, \eta^2 = \mathbf{0.44}$ | n.s. |
| Food Stimulus Investigation [sec] | $F_{(1,15)} = 0.67, p = 0.43, \eta^2 = 0.043$ | $F_{(1,15)} = 0.043, p = 0.84, \eta^2 = 0.003$ | n.s. |
| Total Investigation [sec] | $F_{(1,15)} = 0.10, p = 0.75, \eta^2 = 0.007$ | $F_{(1,15)} = \mathbf{8.99}, p = \mathbf{0.009}, \eta^2 = \mathbf{0.38}$ | n.s. |
| Total Distance [m] | $F_{(1,15)} = 1.06, p = 0.32, \eta^2 = 0.066$ | $F_{(1,15)} = 0.03, p = 0.87, \eta^2 = 0.002$ | n.s. |
| Middle Chamber Entries [#] | $F_{(1,15)} = 0.08, p = 0.79, \eta^2 = 0.005$ | $F_{(1,15)} = 2.33, p = 0.15, \eta^2 = 0.13$ | n.s. |

**Fig 7.**
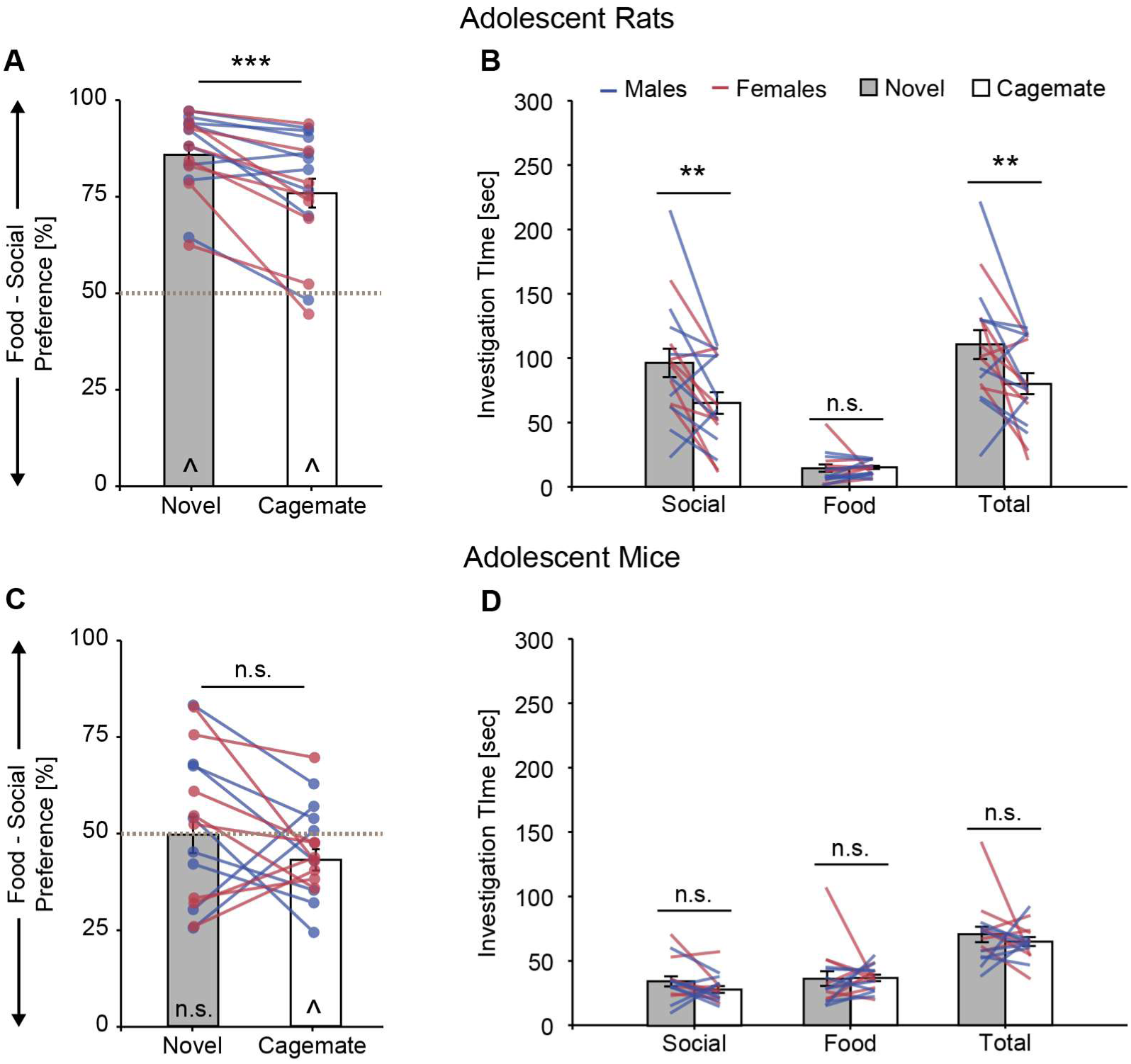
Experiments 4a and 4b. Adolescent rats exhibited a robust preference for the social stimulus, which was attenuated when tested with their cagemate compared to when tested with a novel social stimulus (**A**). Adolescent rats spent more time investigating the social stimulus when it novel than when it was their cagemate; food investigation was low and similar across conditions (**B**). Adolescent mice exhibited a preference for the food stimulus when the social stimulus was their cagemate (**C, left**), but there were no significant changes in preference (**C**) or investigation times (**D**) between social stimulus conditions. Bar graphs (mean ± SEM) collapsed across sex; ** *p* < 0.01, *** *p* < 0.001, mixed-model ANOVA; ^ p < 0.05, one-sample t-Test from 50% (gray dashed line); n.s. = not significant; rats: n = 9 males, n = 8 females; mice: n = 8 males, n = 8 females.

Adolescent rats spent significantly more time investigating the social stimulus when it was novel compared to when it was their cagemate which resulted in a significantly greater total stimulus investigation time when the social stimulus was novel; investigation of the food stimulus was low and similar across conditions (**Table 8, Fig 7B**). There were no sex differences in social, food, or total stimulus investigation times in adolescent rats (**Table 8, Fig 7B**).

Neither sex nor social saliency condition affected locomotor activity as measured by total distance traveled or middle chamber entries in adolescent rats (**Tables 8, 9**).

**Table 9.** Activity measures for Experiments 4a and 4b. Data shown as mean ± SEM; ^1^main effect of sex; see text for details and Tables 8 and 10 for corresponding ANOVA statistics.

|  |  | Cagemate | Novel |
| --- | --- | --- | --- |
| *Total Distance Traveled [m]* |  |  |  |
| Exp 4a:<br>Adolescent Rats | Males | 27.0 ± 2.05 | 24.6 ± 1.64 |
| | Females | 28.3 ± 4.70 | 30.0 ± 2.05 |
| Exp 4b:<br>Adolescent Mice<sup>1</sup> | Males | 18.6 ± 1.36 | 18.2 ± 1.94 |
| | Females | 23.0 ± 0.83 | 24.8 ± 1.73 |
| *Middle Chamber Entries [#]* |  |  |  |
| Exp 4a:<br>Adolescent Rats | Males | 28.7 ± 2.11 | 23.1 ± 1.59 |
| | Females | 25.4 ± 4.64 | 24.5 ± 2.58 |
| Exp 4b:<br>Adolescent Mice<sup>1</sup> | Males | 18.9 ± 2.09 | 21.1 ± 2.56 |
| | Females | 25.9 ± 1.36 | 30.9 ± 4.59 |

### 3.7 Experiment 4b: Stimulus preference was altered by social familiarity in adolescent mice

Adolescent mice did not show a significant within-subjects change in stimulus preference as a result of social saliency (**Fig 7C, Table 10**), and stimulus preference was similar between males and females. However, while adolescent mice did not have a stimulus preference when the social stimulus was novel (*t*_(15)_ = 0.099, *p* = 0.92, *d* = 0.025; **Fig 7C**, right), they exhibited a significant preference for the food stimulus when the social stimulus was their cagemate (*t*_(15)_ = 2.48, *p* = 0.026, *d* = 0.62; **Fig 7C**, left).

**Table 10.**
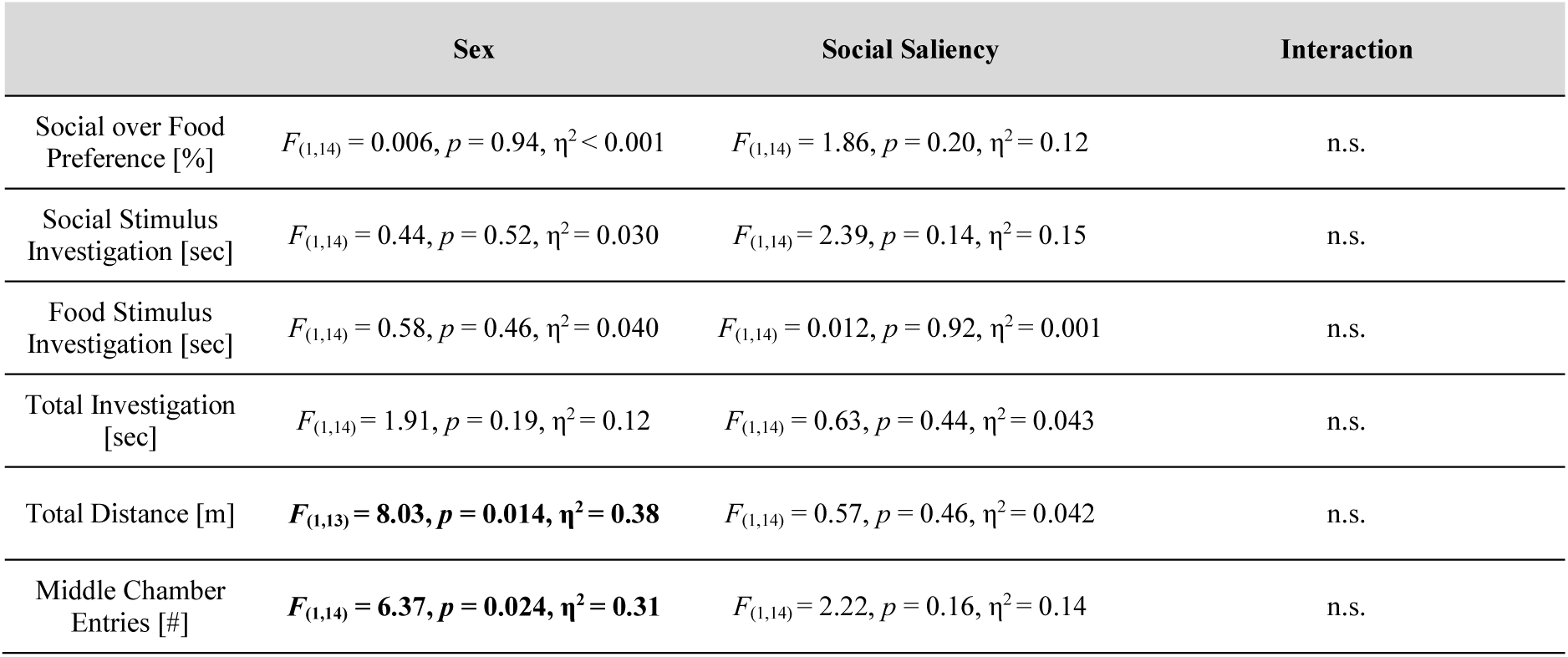
ANOVA statistics and partial eta squared (η^2^) effect sizes for Experiment 4b: Stimulus preference was altered by social familiarity in adolescent mice. Significant effects shown in **bold**, n.s. = none significant.

|  | Sex | Social Saliency | Interaction |
| --- | --- | --- | --- |
| Social over Food Preference [%] | $F_{(1,14)} = 0.006, p = 0.94, \eta^2 < 0.001$ | $F_{(1,14)} = 1.86, p = 0.20, \eta^2 = 0.12$ | n.s. |
| Social Stimulus Investigation [sec] | $F_{(1,14)} = 0.44, p = 0.52, \eta^2 = 0.030$ | $F_{(1,14)} = 2.39, p = 0.14, \eta^2 = 0.15$ | n.s. |
| Food Stimulus Investigation [sec] | $F_{(1,14)} = 0.58, p = 0.46, \eta^2 = 0.040$ | $F_{(1,14)} = 0.012, p = 0.92, \eta^2 = 0.001$ | n.s. |
| Total Investigation [sec] | $F_{(1,14)} = 1.91, p = 0.19, \eta^2 = 0.12$ | $F_{(1,14)} = 0.63, p = 0.44, \eta^2 = 0.043$ | n.s. |
| Total Distance [m] | $F_{(1,13)} = \mathbf{8.03}, p = \mathbf{0.014}, \eta^2 = \mathbf{0.38}$ | $F_{(1,14)} = 0.57, p = 0.46, \eta^2 = 0.042$ | n.s. |
| Middle Chamber Entries [#] | $F_{(1,14)} = \mathbf{6.37}, p = \mathbf{0.024}, \eta^2 = \mathbf{0.31}$ | $F_{(1,14)} = 2.22, p = 0.16, \eta^2 = 0.14$ | n.s. |

There was no effect of sex nor social saliency condition on social, food, or total investigation times in adolescent mice (**Fig 7D, Table 10**).

Locomotor activity was higher in adolescent female mice compared to adolescent male mice as measured by total distance traveled and middle chamber entries, but was unaffected by social saliency condition (**Tables 9, 10**).

### 3.8 Experiment 5a: Adolescent mice preferred to investigate a novel social stimulus over an empty corral

Adolescent mice spent significantly more time investigating the corralled social stimulus compared to the empty corral (*F*_(1,6)_ = 7.10, *p* = 0.037, η^2^ = 0.54; **Fig 8A**). There were no effects of sex on investigation times (*F*_(1,6)_ = 0.38, *p* = 0.56, η^2^ = 0.060; **Fig 8A**) or stimulus preference (*t*_(6)_ = 0.65, *p* = 0.54, *d* = 0.46; **Fig 8B**). Adolescent mice exhibited a significant preference for the social stimulus (*t*_(7)_ = 2.91, *p* = 0.023, *d* = 1.03; **Fig 8B**).

**Fig 8.**
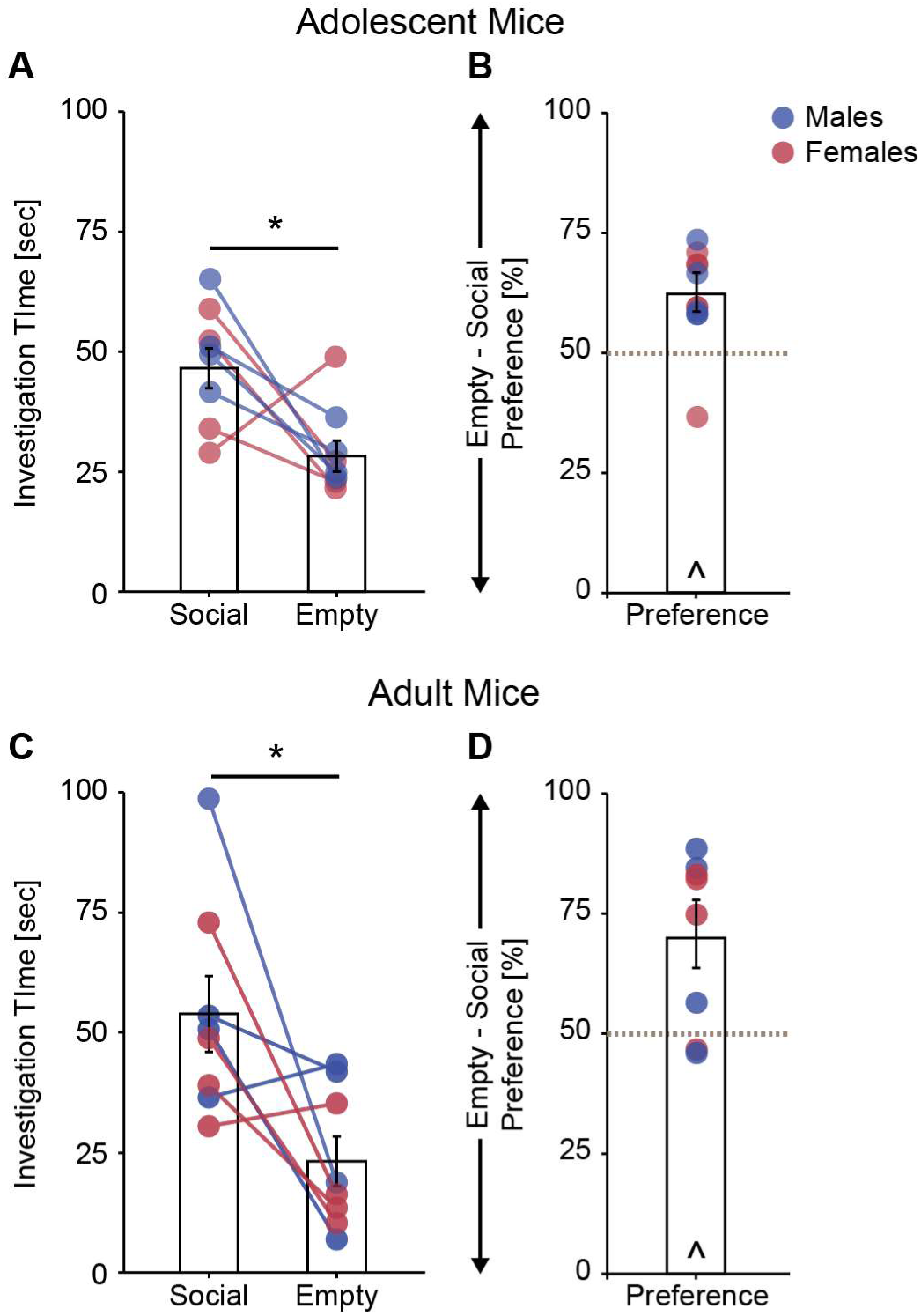
Experiments 5a and 5b. Adolescent mice spent more time investigating a corral containing a novel social stimulus than an empty corral (**A**), and as a result showed a significant preference for the corral containing the social stimulus (**B**). Adult mice spent more time investigating a corral containing a novel social stimulus than an empty corral (**C**), and as a result showed a significant preference for the corral containing the social stimulus (**D**). Bar graphs (mean ± SEM) collapsed across sex; * *p* < 0.05 mixed-model ANOVA; ^ p < 0.05, one-sample t-Test from 50% (gray dashed line); adolescents: n = 4 males, n = 4 females; adults: n = 4 males, n = 4 females).

There were no sex differences in middle chamber entries (males: 32 ± 2.3, females: 37 ± 3.1; *t*_(6)_ = 1.31, *p* = 0.24, *d* = 0.92), or total distance traveled (males: 29.1 ± 1.3 m, females: 25.4 ± 2.2 m; *t*_(6)_ = 1.56, *p* = 0.17, *d* = 1.10) in adolescent mice.

### 3.9 Experiment 5b: Adult mice preferred to investigate a novel social stimulus over an empty corral

Adult mice spent significantly more time investigating the corralled social stimulus compared to the empty corral (*F*_(1,6)_ = 7.00, *p* = 0.038, η^2^ = 0.54; **Fig 8C**). There were no effects of sex on investigation times (*F*_(1,6)_ = 1.95, *p* = 0.21, η^2^ = 0.25; **Fig 8C**) or stimulus preference (*t*_(6)_ = 0.21, *p* = 0.84, *d* = 0.15; **Fig 8D**). Adult mice exhibited a significant preference for the social stimulus (*t*_(7)_ = 3.18, *p* = 0.016, *d* = 1.12; **Fig 8D**).

There were no sex differences in middle chamber entries (males: 17.8 ± 2.7, females: 24.5 ± 5.3; *t*_(6)_ = 1.13, *p* = 0.30, *d* = 0.180), or total distance traveled (males: 15.8 ± 4.7 m, females: 20.2 ± 1.58 m; *t*_(6)_ = 1.53, *p* = 0.18, *d* = 0.46) in adult mice.

## 4 DISCUSSION

Here we characterized a behavioral paradigm designed to test the competition between the choice to seek social interaction versus food, and assessed how this competition was modulated by internal cues, external cues, sex, age, and rodent model. We demonstrated that changes in stimulus preference in response to our cue manipulations (i.e., food deprivation, social isolation, social salience; **Fig 9C, F**) were similar across cohorts (i.e., sex, age, and/or rodent model), but that baseline stimulus preference and investigation times varied greatly between Wistar rats and C57BL/6J mice. Specifically, Wistar rats were generally more social-preferring and C57BL/6J mice were generally more food-preferring (**Fig 9B, E**). Further, especially in Wistar rats, the degree of food deprivation-induced changes in investigation patterns appeared greater in adolescents compared to adults (**Fig 9A-C**). Given these cohort differences, our results highlight the importance of taking experimental population (i.e., age, rodent model) into account when using the Social versus Food Preference test in future experiments.

**Fig 9.**
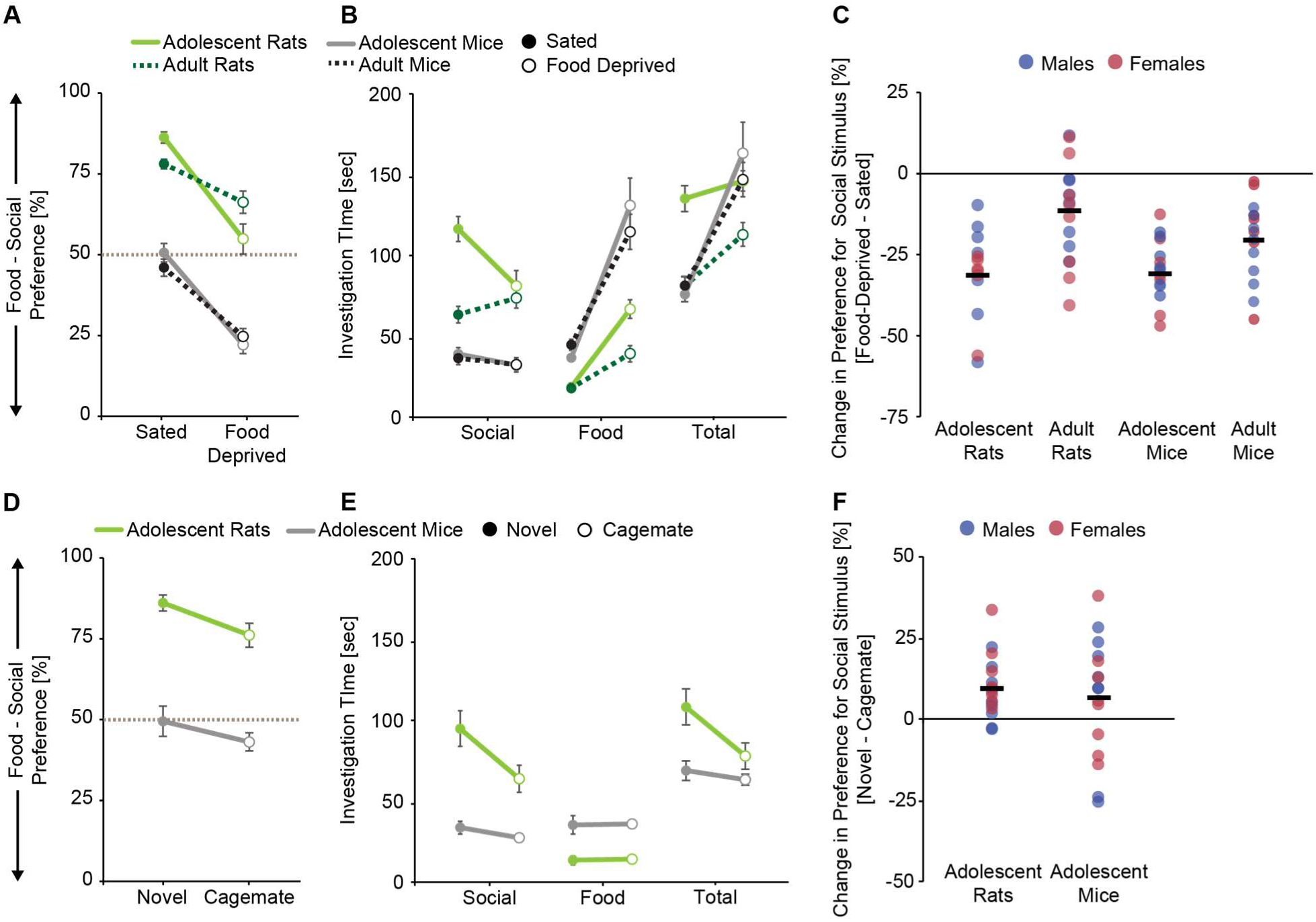
Age and species differences in stimulus preference and investigation times in the Social versus Food Preference Test. Data has been replotted to visually compare the results between Experiments 1a, 1b, 3a, and 3b (**A-C;** data collapsed across sex and housing condition) and between Experiments 4a and 4b (**D-F**; data collapsed across sex); **A, B, D, E**: data shown as mean ± SEM; **C, F**: black bars = mean.

### 4.1 The effects of internal cue manipulations on social versus food preference

To begin characterizing the Social versus Food Preference Test, we first determined how altering the internal motivational states of subjects by exposing them to acute social isolation and/or acute food deprivation affected stimulus preference and stimulus investigation times (Experiments 1, 3). In agreement with our predictions, adolescent and adult rats and mice of both sexes showed a significant reduction in their social over food preference scores in response to acute food deprivation (**Fig 9C**). This change was primarily driven by a significant increase in investigation of the food stimulus in response to food deprivation in all four cohorts (**Fig 9B**). This increased interest in the food stimulus corresponded with our proxy physiological and behavioral measures of hunger; acute food deprivation significantly decreased body weight and increased post-test food consumption compared to sated conditions in all four cohorts (**Table 11**).

**Table 11.**
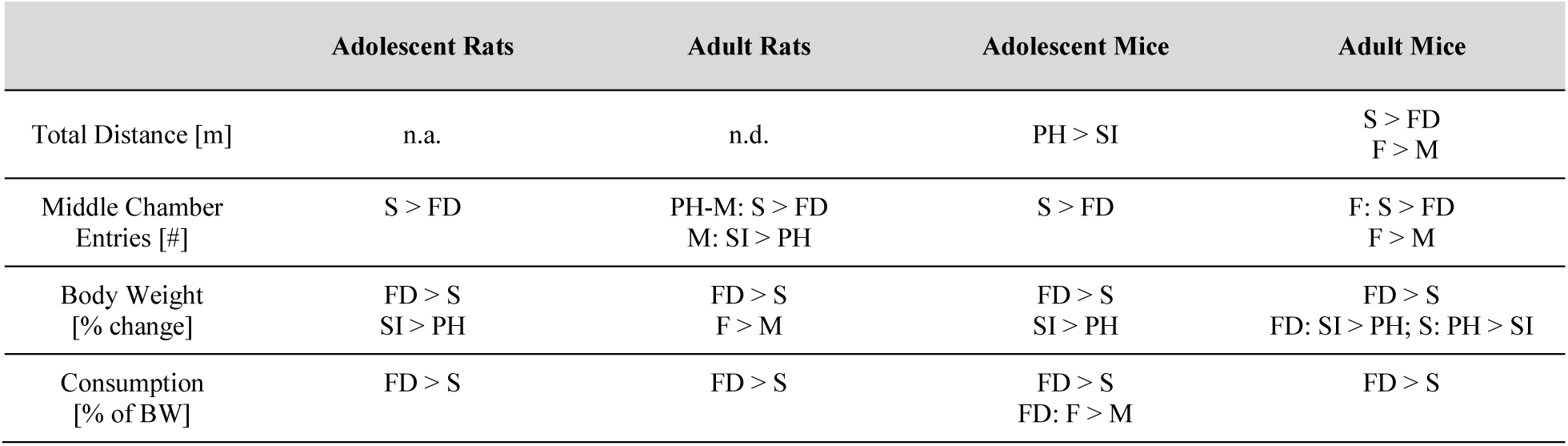
Comparing the results between Experiments 1a, 1b, 3a, 3b for locomotor activity, body weight, and food consumption measures; n.a. = not analyzed; n.d. = no differences between groups or conditions; M = males; F = females; S = sated; FD = food-deprived; PH = pair-housed; SI = socially isolated.

|  | Adolescent Rats | Adult Rats | Adolescent Mice | Adult Mice |
| --- | --- | --- | --- | --- |
| Total Distance [m] | n.a. | n.d. | PH > SI | S > FD<br>F > M |
| Middle Chamber Entries [#] | S > FD | PH-M: S > FD<br>M: SI > PH | S > FD | F: S > FD<br>F > M |
| Body Weight [% change] | FD > S<br>SI > PH | FD > S<br>F > M | FD > S<br>SI > PH | FD > S<br>FD: SI > PH; S: PH > SI |
| Consumption [% of BW] | FD > S | FD > S | FD > S<br>FD: F > M | FD > S |

#### 4.1.1 Age differences in food deprivation-induced changes in stimulus preference and investigation times

While food deprivation decreased social over food preference scores in both adolescents and adults, the degree of change was different between the age groups (i.e., slope of lines in **Fig 9A**). Specifically, the mean difference in preference scores between sated and food-deprived conditions was greater in adolescents compared to adults for both rats and mice (**Fig 9C**). This age difference was partly due to differences in food deprivation-induced changes in social stimulus investigation in adolescents compared to adults. Adolescent rats and mice significantly reduced their investigation of the social stimulus when food-deprived compared to when sated, and this effect size was larger in rats (**Tables 1, 6**; **Fig 9B**). In contrast, adult rats and mice maintained a similar level of social stimulus investigation under both sated and food-deprived conditions (**Tables 3, 7; Fig 9B**). Further, while all cohorts significantly increased their investigation of the food stimulus under food-deprived compared to sated conditions (**Fig 9B**), the effect size was greater in adolescents than adults in both rats and mice (**Tables 1, 3, 6, 7**). These results suggest that adolescents may have been more affected by the food deprivation manipulation when compared to adults. In support, under food-deprived conditions adolescents lost, on average, a greater percent of their body weight, and consumed, on average, more food relative to their body weight compared to adults (**Table 2**).

In rats, we also observed age differences in the absolute amount of time subjects spent investigating the stimuli, and this varied by hunger state. Under sated conditions, adolescent rats spent, on average, 183 % more time than adults investigating the social stimulus, and under food-deprived conditions adolescents spent, on average, 171 % more time than adult rats investigating the food stimulus (**Fig 9B**). Interestingly, investigation of the food stimulus was similar between adolescent rats and adult rats under sated conditions suggesting similar baseline levels in food motivation between the ages, while food deprivation reduced the time adolescent rats spent investigating the social stimulus down to adult rat levels (**Fig 9B**). These age differences are in alignment with increased reward-seeking behavior and motivation for a variety of rewards in adolescents compared to adults, characteristics believed to be evolutionarily important for typical development [for review see: 15, 16]. Since this adolescent-specific peak in reward-seeking and motivated behaviors has been observed across species, including mice, it was surprising that we observed similar amounts of stimulus investigation in adolescent versus adult mice (**Fig 9B**). One reason could be the age of testing for the adolescent mice. Indeed, an earlier study reporting greater levels of social investigation by adolescent compared to adult mice [17], tested adolescent mice at a younger age (30-32 days-old) than we did in the current study (37-44 days-old). Thus, although we attempted to equilibrate the adolescent time-point between species in our experiments [13], differences may exist between rats and mice in the developmental time course of increased reward-seeking behaviors.

#### 4.1.2 Species differences in baseline stimulus preference and investigation times

While social over food preference scores in response to food deprivation were similarly decreased in Wistar rats and C57BL/6J mice, baseline preference scores were vastly different between these two rodent models. Wistar rats exhibited a strong social preference when tested under sated conditions and this was abolished (in adolescents) or attenuated (in adults) by food deprivation (**Fig 9A**). In contrast, adolescent and adult C57BL/6J mice had no stimulus preference when tested under sated conditions and had a strong food preference when food-deprived (**Fig 9A**). This partially matched a study where adult male mice preferred a food stimulus over a social stimulus when food-deprived, but preferred the social stimulus over the food stimulus when sated [3]. Methodological differences between the two studies may account for the differing results under sated conditions, specifically, the length of social isolation (18 hrs in our study vs. > 2.5 weeks in [3]), type of social stimulus (age-, and sex-matched in our study vs. receptive female or juvenile male in [3]), food presentation (corralled in our study vs. available for consumption in [3]), and/or chamber design (3 chambers which provided a neutral zone in our study vs. 2 chambers which forced choice in [3]).

The distinct difference in stimulus preference between Wistar rats and C57BL/6J mice, which was also observed in our experiments investigating the role of social salience (**Fig 9D, Section 4.2.2** below), was driven by differences in the time spent investigating both the social and the food stimuli. Collapsed across experiment (Experiments 1, 3, and 4), age (adolescents, adults), sex (males, females), and manipulation (hunger, housing, and social salience conditions), Wistar rats spent on average 243 % more time investigating the social stimulus than C57BL/6J mice, and C57BL/6J mice spent on average 240 % more time investigating the food stimulus than Wistar rats (**Fig 9B, E**). Thus, our results support the hypothesis that rats and mice differ in their baseline motivation to seek social interaction (higher in rats) and to seek food (higher in mice). This may be due to differences in the evolutionary or ecological history of these rodent species, and corresponds with multiple lines of evidence suggesting that, socially, rats are more agonistic and mice are more antagonistic [for reviews see: 18, 19]. Further, our results are in agreement with a prior study that compared 6-8 week-old male Sprague-Dawley rats and C57BL/6N mice, and found that rats were more likely than mice to develop a social conditioned place preference and that rats spent longer interacting with the social stimulus than mice during the initial conditioning sessions [20].

Differences in the choice to seek social interaction versus food could also be influenced by species differences in metabolism [21, 22]. Indeed, although the length of food deprivation used in the current study was longer for rats than mice (24 hrs versus 18 hrs), mice lost, on average, a greater percent of their body weight in response to food deprivation than rats suggesting mice have higher metabolic demands (**Table 2**). Lastly, it is important to recognize that only one strain of each species was compared in the present study, and that within-species strain differences exist in other social [e.g., 9, 23, 24-27] and food-related behavioral assays [e.g., 28, 29-32]. Thus, future experiments utilizing additional strains of each species are needed to determine if the results from the present study are reflective of each species as a whole.

#### 4.1.3 Males and females exhibited similar stimulus preferences and investigation times

No sex differences in stimulus preference or stimulus investigation times were observed in 8 of the 9 experiments in the current study. The exception was Experiment 2, where adolescent male rats investigated the social stimulus more than adolescent female rats while investigation of the food stimulus was similar between the sexes, resulting in a higher social over food preference score in males compared to females (especially under food-deprived conditions). This was in contrast to Experiments 1a and 4a where adolescent male and female rats showed similar levels of social stimulus investigation. It could be that males were less affected by the food deprivation manipulation than females in Experiment 2 (i.e., males did not lose as high of a percent of their body weight as females; males: 8.2 %, females: 11.1 %), or than males in Experiment 1a (10.3 %). Additionally, across conditions, males in Experiment 2 investigated the social stimulus more than males in Experiment 1a or 4a, while females investigated the social stimulus similar amounts across experiments. The increased investigation in males in Experiment 2 could be due to differences in housing conditions between the experiments (i.e., acute social isolation and/or pair-housed in Experiments 1a and 4a, single-housed for the entire duration of Experiment 2), which is an important question for future research (see **Section 4.1.4**). Alternatively, we have previously observed individual and/or cohort variability in social investigation by young rats in other behavioral paradigms [8, 10, 33, 34]. Thus, despite some cohort variability, as a whole, males and females exhibited similar patterns of stimulus preference and levels of stimulus investigation in the Social versus Food Preference Test under all examined conditions.

#### 4.1.4 Social isolation did not affect stimulus investigation patterns, but did potentiate food deprivation-induced weight loss

Contrary to our prediction, acute social isolation did not affect stimulus preference or stimulus investigation in any cohort (**Fig 9A-C**). Our prediction was based on prior work reporting increased social investigation by rats and mice following short-term (3-24 hr) social isolation [e.g., 10, 35, 36, 37]. Additionally, a recent meta-analysis found that independent of age or sex, rats and mice consume more food when socially isolated compared to when socially housed [38] suggesting that social isolation may also increase food-directed motivation. It should be noted, however, that the minimum amount of social isolation examined in this meta-analysis (48 hr) was at least twice that used in our current study (24 hrs in rats, 18 hrs in mice). A social isolation-induced increase in food-directed motivation may be an adaptive response since socially isolated subjects are unable to engage in huddling, a behavior commonly expressed by rats and mice [39-41] and which is thought to serve a thermoregulatory purpose [42]. Thus, these long-term socially isolated subjects may have had increased metabolic demands that manifested as increased food-directed motivation. The shorter length of social isolation used in our current study may not have been long enough to induce measurable changes in social- or food-directed motivation and subsequent behavior on the Social versus Food Preference Test. In support, an earlier study found that sated adult male mice that had been isolated for at least 2.5 weeks preferred a social stimulus (receptive female or juvenile male) over a food stimulus (standard laboratory chow) [3]. Further, we equated the absolute duration of food deprivation and social isolation in our experiments (i.e., 24 hrs for both manipulations in rats, 18 hrs for both manipulations in mice), but this may not elicit equal changes in the motivation to seek social interaction compared to the motivation to seek food. Future studies could determine what length of social isolation would induce changes in behavior within this paradigm.

Social isolation did consistently alter one parameter examined in the current set of experiments: percent change in body weight (**Table 11**). Specifically, adolescent rats and adolescent and adult mice lost more weight under food deprivation when they were socially isolated compared to when they were pair-housed. Social isolation-induced changes in body weight were not observed in adult rats, which may reflect differences in metabolic demands corresponding to body size [22] or within-species age differences in spontaneous activity [43]. One likely explanation for increased body weight loss is the inability of socially isolated subjects to engage in huddling [39, 40, 42]. Thus, socially isolated subjects may have expended more energy for thermoregulation processes than pair-housed subjects, and in the absence of food were unable to compensate with an increase in energy intake. In future studies, conducting homecage behavioral observations and monitoring food consumption during the social isolation period could provide evidence to support this hypothesis.

### 4.2 The effects of external cue manipulations on social versus food preference

Since behavior is influenced both by internal and external cues [1, 2], we next sought to determine how behavior in the Social versus Food Preference Test would be affected by manipulations of external cues. Since adolescents showed greater changes in behavior compared to adults in our manipulations of internal cues (i.e., Experiments 1, 3; see **Section 4.1.1**), we chose to only use adolescents for these experiments. First, we examined whether the time-of-testing (i.e., light phase versus dark phase) influenced stimulus investigation or preference in adolescent male and female rats (Experiment 2), and next we examined whether manipulating the saliency of the social stimulus (i.e., novel versus familiar) would alter social versus food preference in sated adolescent rats and mice of both sexes (Experiment 4).

#### 4.2.1 The time-of-testing did not influence stimulus investigation patterns

Laboratory rats and mice are nocturnal species that exhibit increased locomotor activity during the dark phase compared to the light phase [7], and this is true across age and sex [44]. When tested in a home environment, increased social interactions at the transition from the light to the dark phase correlates with this increased locomotor activity [36], and the expression of a variety of other spontaneous social behaviors in the home environment also varies across the light cycle [39]. Similarly, patterns of food intake in a home environment vary with light cycle with males and females consuming more food during the dark phase (especially during the early dark phase) compared to the light phase [45-47]. Thus, we predicted that the relative preference of socially isolated adolescent rats to investigate a social stimulus versus a food stimulus would be similar between the dark phase and light phase, but that overall interest in both the social and food stimuli and/or general locomotor activity may be higher during the dark phase compared to the light phase. In fact, we did not observe any effects of light phase on stimulus preference, stimulus investigation times, or locomotor activity in the Social versus Food Preference Test. This is consistent with prior studies where socially isolated adult male rats showed similar levels of social investigation [35, 48], and exhibited similar latencies to locate and begin eating a palatable food as well as consumed similar amounts of that palatable food [49] in the light phase versus the dark phase. Together, this suggests that while baseline activity in the home environment reliably varies across the light cycle, activity elicited in a behavioral test may be less affected by the light cycle. This supports the future use of the Social versus Food Preference Test during the light phase which can alleviate practical difficulties associated with dark phase testing [50].

#### 4.2.2 Social saliency modulated stimulus preference

Given the previously described preference of rats and mice for social novelty [8-10, 23, 24, 51], and the increased novelty-seeking behavior displayed by adolescents across species [for review see: 15, 16], we predicted that social novelty, as opposed to social familiarity, would bias preference more towards the social stimulus in the Social versus Food Preference Test. In general, our prediction was confirmed; adolescent rats displayed a stronger preference for the social stimulus when the social stimulus was a novel conspecific compared to when it was their cagemate, and while adolescent mice did not have a stimulus preference when the social stimulus was a novel conspecific they preferred the food stimulus when the social stimulus was their cagemate (**Fig 9D**). These results further reinforce our working hypothesis that while the baseline balance between social motivation and food motivation differs between rats and mice (see **Section 4.1.2**), how rats and mice respond to experimental manipulations of these motivations is similar.

### 4.3 Despite a lack of social over food preference, mice prefer a social stimulus over an empty corral

Prior studies on general sociability have consistently shown that C57BL/6J mice prefer a social stimulus over an object [23, 24] or empty chamber [9]. In agreement with these prior studies, we found that adolescent and adult male and female C57BL/6J mice showed a significant preference for a corralled social stimulus versus an empty corral. When considered with the results from Experiments 3 and 4b this suggests that, in C57BL/6J mice tested under stated conditions, the motivation to seek food is similar to the motivation to seek social contact, and both are greater than the motivation to investigate an object. However, it should be noted this refers to motivation as measured by choice in a passive investigation paradigm. In an operant two-choice lever-pressing paradigm, sated adult male C57BL/6J and BTBR T+tf/J mice had greater motivation for a palatable food reward (sweetened evaporated milk) compared to a social reward (sex- and age-matched stimulus mouse) [32]. Whether the difference in the balance between social motivation and food motivation between this prior study and the current study was due to differences in the nature of the task (operant in [32] versus passive investigation in current study) or the value of the food stimulus (palatable in [32] versus standard laboratory chow in current study) could be addressed in future studies by using a palatable food stimulus in the Social versus Food Preference Test.

### 4.4 The Social versus Food Preference Test is a flexible behavioral paradigm for studying competing motivations

The present series of experiments highlight the flexible nature of the Social versus Food Preference Test, and support its use in future studies of motivated behavioral choice and interrogations of the underlying pheripheral and central systems. A key benefit of this behavioral test is the ability to assess both absolute (i.e., investigation time) and relative (i.e., preference) interest to investigate two opposing stimuli, giving multiple readouts for interpreting how manipulations of test parameters or neural systems can affect behavior. Further, the ease of manipulating internal motivational states (e.g., varying the length of food deprivation) and/or the salience or reward-value of the external stimuli (e.g., novelty, palatability, stimulus devaluation) to alter behavior, will allow experimenters to titer stimulus preference or investigation times. When combined with careful consideration of experimental population, this ability will be key in tailoring experiments for specific hypotheses and preventing floor or ceiling effects (e.g., prediction for decreased social preference: select sated Wistar rats; prediction for decreased food preference: select food-deprived C57BL/6J mice; uncertain direction of prediction: select sated C57BL/6J mice or food-deprived Wistar rats). Lastly, while sex differences in the current set of behavioral experiments were minimal, we would encourage the continued use of both sexes in future investigations of the neural substrates underlying social versus food preference [52, 53]. This is critical because an absence of sex differences in behavior could still be caused by sex differences in its neural underpinnings [54], and we and others have demonstrated that the neural mechanisms underlying motivated social behavior can differ in males and females [34, 55-60].

## 5 CONCLUSIONS

Here we established the Social versus Food Preference Test to examine the competition between the choice to seek social interaction versus the choice to seek food. First, we characterized how this competition was modulated by internal cues (i.e., acute food deprivation and/or acute social isolation) and assessed whether these manipulations would produce similar changes in behavior between the sexes (males, females), across the lifespan (adolescents, adults), and between commonly used laboratory rodent models (Wistar rats, C57BL/6J mice). We found that behavior in this test was similar between the sexes and unaffected by social isolation, but highly influenced by food deprivation (i.e., biased preference more towards the food stimulus) and that this effect size was larger in adolescents than adults. Most strikingly, we observed a robust baseline difference in preference between Wistar rats and C57BL/6J mice: Wistar rats were generally more social-preferring and C57BL/6J mice were generally more food-preferring. Next, we determined whether the competition between the choice to seek social interaction versus the choice to seek food could be modulated by external cues and found that behavior was similar in the light phase versus the dark phase in adolescent male and female Wistar rats, but that behavior was altered by changing the salience of the social stimulus (i.e., social novelty biased preference more towards the social stimulus) in adolescent Wistar rats and C57BL/6J mice of both sexes. Together, our experiments confirm that the Social versus Food Preference Test is a flexible behavioral paradigm suitable for future interrogations of the peripheral and central systems that can coordinate the expression of stimulus preference related to multiple motivated behaviors.

## ACKNOWLEDGMENTS

We would like to thank members of the Veenema Lab for critical review of prior drafts of this manuscript, and Natasha M. Méndez Albelo, Suhana S. Posani, and Sang Y. Yang for technical assistance.

## AUTHORS’ CONTRIBUTIONS

CJR and AHV conceived the idea and designed the experiments; CJR, LAB, and AQC conducted the experiments; CJR analyzed the data and wrote the first draft of the manuscript; AHV supervised the project; all authors discussed the results and commented on the manuscript.

## REFERENCES

1. Mogenson, G.J., The neurobiology of behavior: an introduction. 1977, New York, NY: L. Erlbaum Associates. 334.

2. Berridge, K.C. 2004. Motivation concepts in behavioral neuroscience. Physiol Behav. 81(2), 179–209. DOI: 10.1016/j.physbeh.2004.02.004

3. Burnett, C.J., Li, C., Webber, E., Tsaousidou, E., Xue, S.Y., Bruning, J.C., & Krashes, M.J. 2016. Hunger-driven motivational state competition. Neuron. 92(1), 187–201. DOI: 10.1016/j.neuron.2016.08.032

4. Padilla, S.L., Qiu, J., Soden, M.E., Sanz, E., Nestor, C.C., Barker, F.D., … Palmiter, R.D. 2016. Agouti-related peptide neural circuits mediate adaptive behaviors in the starved state. Nat Neurosci. 19(5), 734–741. DOI: 10.1038/nn.4274

5. Jennings, J.H., Kim, C.K., Marshel, J.H., Raffiee, M., Ye, L., Quirin, S., … Deisseroth, K. 2019. Interacting neural ensembles in orbitofrontal cortex for social and feeding behaviour. Nature. 565(7741), 645–649. DOI: 10.1038/s41586-018-0866-8

6. Yoest, K.E., Cummings, J.A., & Becker, J.B. 2019. Ovarian hormones mediate changes in adaptive choice and motivation in female rats. Front Behav Neurosci. 13, 250. DOI: 10.3389/fnbeh.2019.00250

7. Refinetti, R. 2006. Variability of diurnality in laboratory rodents. J Comp Physiol A Neuroethol Sens Neural Behav Physiol. 192(7), 701–14. DOI: 10.1007/s00359-006-0093-x

8. Smith, C.J., Wilkins, K.B., Mogavero, J.N., & Veenema, A.H. 2015. Social novelty investigation in the juvenile rat: Modulation by the mu-opioid system. J Neuroendocrinol. 27(10), 752–64. DOI: 10.1111/jne.12301

9. Moy, S.S., Nadler, J.J., Perez, A., Barbaro, R.P., Johns, J.M., Magnuson, T.R., … Crawley, J.N. 2004. Sociability and preference for social novelty in five inbred strains: an approach to assess autistic-like behavior in mice. Genes Brain Behav. 3(5), 287–302. DOI: 10.1111/j.1601-1848.2004.00076.x

10. Smith, C.J.W., Wilkins, K.B., Li, S., Tulimieri, M.T., & Veenema, A.H. 2018. Nucleus accumbens mu opioid receptors regulate context-specific social preferences in the juvenile rat. Psychoneuroendocrinology. 89, 59–68. DOI: 10.1016/j.psyneuen.2017.12.017

11. Bull, L.S. & Pitts, G.C. 1971. Gastric capacity and energy absorption in the force-fed rat. J Nutr. 101(5), 593–6. DOI: 10.1093/jn/101.5.593

12. Asarian, L. & Geary, N. 2013. Sex differences in the physiology of eating. Am J Physiol Regul Integr Comp Physiol. 305(11), R1215–67. DOI: 10.1152/ajpregu.00446.2012

13. Schneider, M. 2013. Adolescence as a vulnerable period to alter rodent behavior. Cell Tissue Res. 354(1), 99–106. DOI: 10.1007/s00441-013-1581-2

14. Weissgerber, T.L., Milic, N.M., Winham, S.J., & Garovic, V.D. 2015. Beyond bar and line graphs: time for a new data presentation paradigm. PLoS Biol. 13(4), e1002128. DOI: 10.1371/journal.pbio.1002128

15. Spear, L.P. 2011. Rewards, aversions and affect in adolescence: emerging convergences across laboratory animal and human data. Dev Cogn Neurosci. 1(4), 392–400. DOI: 10.1016/j.dcn.2011.08.001

16. Walker, D.M., Bell, M.R., Flores, C., Gulley, J.M., Willing, J., & Paul, M.J. 2017. Adolescence and reward: Making sense of neural and behavioral changes amid the chaos. J Neurosci. 37(45), 10855–10866. DOI: 10.1523/JNEUROSCI.1834-17.2017

17. Kareem, A.M. & Barnard, C.J. 1982. The importance of kinship and familiarity in social interactions between mice. Animal Behaviour. 30(2), 594–601. DOI: 10.1016/S0003-3472(82)80073-0

18. Kondrakiewicz, K., Kostecki, M., Szadzinska, W., & Knapska, E. 2019. Ecological validity of social interaction tests in rats and mice. Genes Brain Behav. 18(1), e12525. DOI: 10.1111/gbb.12525

19. Ellenbroek, B. & Youn, J. 2016. Rodent models in neuroscience research: is it a rat race? Dis Model Mech. 9(10), 1079–1087. DOI: 10.1242/dmm.026120

20. Kummer, K.K., Hofhansel, L., Barwitz, C.M., Schardl, A., Prast, J.M., Salti, A., … Zernig, G. 2014. Differences in social interaction-vs. cocaine reward in mouse vs. rat. Front Behav Neurosci. 8, 363. DOI: 10.3389/fnbeh.2014.00363

21. Menahan, L.A. & Sobocinski, K.A. 1983. Comparison of carbohydrate and lipid metabolism in mice and rats during fasting. Comp Biochem Physiol B. 74(4), 859–64. DOI: 10.1016/0305-0491(83)90157-8

22. White, C.R. & Kearney, M.R. 2013. Determinants of inter-specific variation in basal metabolic rate. J Comp Physiol B. 183(1), 1–26. DOI: 10.1007/s00360-012-0676-5

23. Moy, S.S., Nadler, J.J., Young, N.B., Perez, A., Holloway, L.P., Barbaro, R.P., … Crawley, J.N. 2007. Mouse behavioral tasks relevant to autism: phenotypes of 10 inbred strains. Behav Brain Res. 176(1), 4–20. DOI: 10.1016/j.bbr.2006.07.030

24. Moy, S.S., Nadler, J.J., Young, N.B., Nonneman, R.J., Segall, S.K., Andrade, G.M., … Magnuson, T.R. 2008. Social approach and repetitive behavior in eleven inbred mouse strains. Behav Brain Res. 191(1), 118–29. DOI: 10.1016/j.bbr.2008.03.015

25. Northcutt, K.V. & Nwankwo, V.C. 2018. Sex differences in juvenile play behavior differ among rat strains. Dev Psychobiol. 60(8), 903–912. DOI: 10.1002/dev.21760

26. Siviy, S.M., Love, N.J., DeCicco, B.M., Giordano, S.B., & Seifert, T.L. 2003. The relative playfulness of juvenile Lewis and Fischer-344 rats. Physiology & Behavior. 80(2-3), 385–394. DOI: 10.1016/j.physbeh.2003.09.002

27. Panksepp, J.B. & Lahvis, G.P. 2007. Social reward among juvenile mice. Genes Brain Behav. 6(7), 661–71. DOI: 10.1111/j.1601-183X.2006.00295.x

28. Clemens, L.E., Jansson, E.K., Portal, E., Riess, O., & Nguyen, H.P. 2014. A behavioral comparison of the common laboratory rat strains Lister Hooded, Lewis, Fischer 344 and Wistar in an automated homecage system. Genes Brain Behav. 13(3), 305–21. DOI: 10.1111/gbb.12093

29. Newmyer, B.A., Whindleton, C.M., Srinivasa, N., Jones, M.K., & Scott, M.M. 2019. Genetic variation affects binge feeding behavior in female inbred mouse strains. Sci Rep. 9(1), 15709. DOI: 10.1038/s41598-019-51874-7

30. Atalayer, D. & Rowland, N.E. 2010. Comparison of C57BL/6 and DBA/2 mice in food motivation and satiety. Physiol Behav. 99(5), 679–83. DOI: 10.1016/j.physbeh.2010.02.001

31. Christensen, C.J., Kohut, S.J., Handler, S., Silberberg, A., & Riley, A.L. 2009. Demand for food and cocaine in Fischer and Lewis rats. Behav Neurosci. 123(1), 165–71. DOI: 10.1037/a0013736

32. Martin, L., Sample, H., Gregg, M., & Wood, C. 2014. Validation of operant social motivation paradigms using BTBR T+tf/J and C57BL/6J inbred mouse strains. Brain Behav. 4(5), 754–64. DOI: 10.1002/brb3.273

33. Reppucci, C.J., Gergely, C.K., Bredewold, R., & Veenema, A.H. 2020. Involvement of orexin/hypocretin in the expression of social play behaviour in juvenile rats. International Journal of Play. 9(1), 108–127. DOI: 10.1080/21594937.2020.1720132

34. Veenema, A.H., Bredewold, R., & de Vries, G.J. 2013. Sex-specific modulation of juvenile social play by vasopressin. Psychoneuroendocrinology. 38(11), 2554–61. DOI: 10.1016/j.psyneuen.2013.06.002

35. Niesink, R.J. & van Ree, J.M. 1982. Short-term isolation increases social interactions of male rats: a parametric analysis. Physiol Behav. 29(5), 819–25. DOI: 10.1016/0031-9384(82)90331-6

36. Panksepp, J.B., Wong, J.C., Kennedy, B.C., & Lahvis, G.P. 2008. Differential entrainment of a social rhythm in adolescent mice. Behav Brain Res. 195(2), 239–45. DOI: 10.1016/j.bbr.2008.09.010

37. Varlinskaya, E.I., Spear, L.P., & Spear, N.E. 1999. Social behavior and social motivation in adolescent rats: role of housing conditions and partner’s activity. Physiol Behav. 67(4), 475–82. DOI: 10.1016/s0031-9384(98)00285-6

38. Schipper, L., Harvey, L., van der Beek, E.M., & van Dijk, G. 2018. Home alone: a systematic review and meta-analysis on the effects of individual housing on body weight, food intake and visceral fat mass in rodents. Obes Rev. 19(5), 614–637. DOI: 10.1111/obr.12663

39. Arakawa, H., Blanchard, D.C., & Blanchard, R.J. 2007. Colony formation of C57BL/6J mice in visible burrow system: identification of eusocial behaviors in a background strain for genetic animal models of autism. Behav Brain Res. 176(1), 27–39. DOI: 10.1016/j.bbr.2006.07.027

40. Barnett, S.A., The rat: A study in behavior. Rev. ed. 1976, Canberra: Australian National University Press. xiv, 318 p.

41. Schweinfurth, M.K. 2020. The social life of Norway rats (Rattus norvegicus). Elife. 9. DOI: 10.7554/eLife.54020

42. Gilbert, C., McCafferty, D., Le Maho, Y., Martrette, J.M., Giroud, S., Blanc, S., & Ancel, A. 2010. One for all and all for one: the energetic benefits of huddling in endotherms. Biol Rev Camb Philos Soc. 85(3), 545–69. DOI: 10.1111/j.1469-185X.2009.00115.x

43. Pereboom, A.C. 1968. Systematic-representative study of spontaneous activity in the rat. Psychol Rep. 22(3), 717–32. DOI: 10.2466/pr0.1968.22.3.717

44. Norton, S., Culver, B., & Mullenix, P. 1975. Development of nocturnal behavior in albino rats. Behav Biol. 15(3), 317–31. DOI: 10.1016/s0091-6773(75)91717-4

45. Balagura, S. & Devenport, L.D. 1970. Feeding patterns of normal and ventromedial hypothalamic lesioned male and female rats. J Comp Physiol Psychol. 71(3), 357–64. DOI: 10.1037/h0029118

46. Siegel, P.S. 1961. Food intake in rat in relation to dark-light cycle. Journal of comparative and physiological Psychology. 54(3), 294-&. DOI: 10.1037/h0044787

47. Siegel, P.S. & Stuckey, H.L. 1947. The diurnal course of water and food intake in the normal mature rat. J Comp Physiol Psychol. 40(5), 365–70. DOI: 10.1037/h0062185

48. Deak, T., Arakawa, H., Bekkedal, M.Y., & Panksepp, J. 2009. Validation of a novel social investigation task that may dissociate social motivation from exploratory activity. Behav Brain Res. 199(2), 326–33. DOI: 10.1016/j.bbr.2008.12.011

49. Tallett, A.J., Blundell, J.E., & Rodgers, R.J. 2009. Night and day: diurnal differences in the behavioural satiety sequence in male rats. Physiol Behav. 97(1), 125–30. DOI: 10.1016/j.physbeh.2009.01.022

50. Yang, M., Weber, M.D., & Crawley, J.N. 2008. Light phase testing of social behaviors: not a problem. Front Neurosci. 2(2), 186–91. DOI: 10.3389/neuro.01.029.2008

51. Eagle, A.L., Fitzpatrick, C.J., & Perrine, S.A. 2013. Single prolonged stress impairs social and object novelty recognition in rats. Behav Brain Res. 256, 591–7. DOI: 10.1016/j.bbr.2013.09.014

52. Cahill, L. 2006. Why sex matters for neuroscience. Nat Rev Neurosci. 7(6), 477–84. DOI: 10.1038/nrn1909

53. Clayton, J.A. & Collins, F.S. 2014. Policy: NIH to balance sex in cell and animal studies. Nature. 509(7500), 282–3. DOI: 10.1038/509282a

54. de Vries, G.J. 2004. Minireview: Sex differences in adult and developing brains: compensation, compensation, compensation. Endocrinology. 145(3), 1063–8. DOI: 10.1210/en.2003-1504

55. Bredewold, R., Smith, C.J.W., Dumais, K.M., & Veenema, A.H. 2014. Sex-specific modulation of juvenile social play behavior by vasopressin and oxytocin depends on social context. Frontiers in Behavioral Neuroscience. 8(216). DOI: 10.3389/Fnbeh.2014.00216

56. Bredewold, R., Schiavo, J.K., van der Hart, M., Verreij, M., & Veenema, A.H. 2015. Dynamic changes in extracellular release of GABA and glutamate in the lateral septum during social play behavior in juvenile rats: Implications for sex-specific regulation of social play behavior. Neuroscience. 307, 117–127. DOI: 10.1016/j.neuroscience.2015.08.052

57. Bredewold, R., Nascimento, N.F., Ro, G.S., Cieslewski, S.E., Reppucci, C.J., & Veenema, A.H. 2018. Involvement of dopamine, but not norepinephrine, in the sex-specific regulation of juvenile socially rewarding behavior by vasopressin. Neuropsychopharmacology. 43(10), 2109. DOI: 10.1038/s41386-018-0100-2

58. Reppucci, C.J., Gergely, C.K., & Veenema, A.H. 2018. Activation patterns of vasopressinergic and oxytocinergic brain regions following social play exposure in juvenile male and female rats. J Neuroendocrinol. DOI: 10.1111/jne.12582

59. Paul, M.J., Terranova, J.I., Probst, C.K., Murray, E.K., Ismail, N.I., & de Vries, G.J. 2014. Sexually dimorphic role for vasopressin in the development of social play. Frontiers in Behavioral Neuroscience. 8. DOI: 10.3389/Fnbeh.2014.00058

60. Northcutt, K.V. & Nguyen, J.M. 2014. Female juvenile play elicits Fos expression in dopaminergic neurons of the VTA. Behav Neurosci. 128(2), 178–86. DOI: 10.1037/a0035964

